# Shared neural codes of recognition memory

**DOI:** 10.1101/2022.12.20.520854

**Authors:** Géza Gergely Ambrus

## Abstract

Recognition memory research has identified several electrophysiological indicators of successful memory retrieval, known as old-new effects. These effects have been observed in different sensory domains using various stimulus types, but little attention has been given to their similarity or distinctiveness and the underlying processes they may share. Here, a data-driven approach was taken to investigate the temporal evolution of shared information content between different memory conditions using openly available EEG data from healthy human participants of both sexes, taken from six experiments. A test dataset involving personally highly familiar and unfamiliar faces was used. The results show that neural signals of recognition memory for face stimuli were highly generalized starting from around 200 ms following stimulus onset. When training was performed on non-face datasets, an early (around 200-300 ms) to late (post-400 ms) differentiation was observed over most regions of interest. Successful cross-classification for non-face stimuli (music and object/scene associations) was most pronounced in late period. Additionally, a striking dissociation was observed between familiar and remembered objects, with shared signals present only in the late window for correctly remembered objects, while cross-classification for familiar objects was successful in the early period as well. These findings suggest that late neural signals of memory retrieval generalize across sensory modalities and stimulus types, and the dissociation between familiar and remembered objects may provide insight into the underlying processes.

## Introduction

Decades of research in the field of recognition memory have identified a number of electrophysiological indicators of successful memory retrieval, known as old-new effects (Rugg and Curran, 2007). These effects have been studied extensively in the context of familiarity and recollection (Brown and Banks, 2015; Wolk et al., 2006; Yonelinas, 2002), and have been observed across a range of sensory domains and stimulus types (Donaldson and Rugg, 1998; Wilding and Rugg, 1997). However, the similarity or distinctiveness of these electrophysiological signals, and the underlying processes that they reflect, have received little attention. Additionally, there have been few attempts to determine whether these findings can be generalized across experiments (Kwon et al., 2022).

In the literature on recognition memory, two key electrophysiological components have been identified: the early mid-frontal effect (FN400) and the late parietal component (LPC). The FN400 typically appears around 300-500 ms after stimulus presentation, predominantly over the frontal regions of the brain, while the LPC is typically observed as a broad positive deflection between 400-800 ms after stimulus onset. Dual process theories of memory retrieval, which posit different neuro-cognitive processes underlying familiarity and recollection, find support in the behavior of these two components under different experimental manipulations. The earlier FN400 has been linked to familiarity, a fast and automatic process associated with “just knowing” a stimulus even in the absence of contextual information. In contrast, the LPC has been associated with recollection, a slower and more controlled process involving the recall of episodic information about previous encounters with the stimulus (Jacoby, 1991; Yonelinas, 2002).

Face perception has often been studied as a model system for investigating the organizational principles of the brain (Young, 2018), but faces are also a unique category of visual stimulus due to their high biological, personal and social importance. This importance is reflected in the prioritization of face detection over other types of visual stimuli (Crouzet, 2010; Morrisey et al., 2019; Visconti di Oleggio Castello and Gobbini, 2015), as well as the preferential processing of familiar faces over unfamiliar ones (Ramon and Gobbini, 2018).

Rapid saccades toward familiar faces have been observed as early as 180 ms after stimulus presentation (Visconti di Oleggio Castello and Gobbini, 2015), and early EEG responses, such as the N170, have been, in cases, shown to be modulated by face familiarity depending on the context of the task (Johnston et al., 2016). However, consistent effects of face familiarity on neural activity do not seem to emerge until after 200 ms post-stimulus. The earliest event-related potential (ERP) component that has been reliably shown to be modulated by face familiarity is the N250 (Huang et al., 2017), a parieto-occipital deflection occurring between 230-350 ms, which is thought to reflect the visual recognition of a known face (Schweinberger and Neumann, 2016). The sustained familiarity effect, peaking between 400-600 ms, is theorized to reflect activity in the “familiarity hub”, where structural information about a face is integrated with additional semantic and affective information (Wiese et al., 2019b).

The interpretation and synthesis of existing findings on the neural basis of recognition memory are limited by the considerable differences in timing and topographical distribution across studies. This means that signals related to memory processes that do not conform to the canonical configurations described above may be overlooked (Dimsdale-Zucker et al., 2022; Kwon et al., 2022). Additionally, the averaged waveforms elicited by the presentation of stimuli are likely to reflect a multitude of temporally overlapping but separate cognitive processes (Campbell et al., 2020), making it difficult to disentangle the underlying neural mechanisms. Therefore, there is a need for further research to better understand the temporal dynamics governing successful memory recall (Staresina and Wimber, 2019).

In this study, instead of focusing on the presence or absence of these signals of successful retrieval in separate experiments, a data-driven approach was taken to probe how shared information content in neural signals between different memory task conditions unfolds over time. A cross-experiment multivariate cross-classification (MVCC, Kaplan et al., 2015) analysis on openly available EEG data from ca. 120 participants of both sexes across six experiments was performed (Dalski et al., 2022a, 2022c, 2022b; Li et al., 2022). The test dataset for this analysis consisted of an experiment involving personally highly familiar and unfamiliar faces (Wiese et al., 2022), and classifier training was conducted using data from two additional face memory experiments (Sommer et al., 2021; Wakeman and Henson, 2015), an experiment presenting short segments of familiar and unfamiliar music (Jagiello et al., 2019), a study investigating remembered and forgotten object-scene associations (Treder et al., 2021), and an object familiarity/recollection study (Dimsdale-Zucker et al., 2022).

In multivariate pattern analysis (MVPA), a machine learning classifier is trained on a set of features (e.g., ERP amplitudes from multiple channels) to find a decision boundary that best separates the categories of interest. This trained classifier is then used to categorize data that was not part of the training set. Higher-than-chance classification indicates that the patterns of data used for training contain information that can be used to successfully classify the test data, indicating that the patterns contain information that separates the categories of interest, and that this information is present in both the training and the test datasets (Grootswagers et al., 2017; Kaplan et al., 2015; King and Dehaene, 2014). One major advantage of MVPA over other methods is its flexibility in terms of what data can be used for training and testing. For example, cross-classification can be performed across different time points, regions of interest, experimental conditions, participants, or datasets from different experiments. Furthermore, pre-existing labels in the training and test datasets can be easily modified, allowing for the evaluation of cross-classification performance across different tasks and domains.

Previous studies have found that neural signals for familiarity with face stimuli generalize across experiments that use different familiarization methods (Dalski et al., 2022a), and that successful cross-classification can be demonstrated even across markedly different task conditions (Dalski et al., 2022c, 2022b). These results suggest that for faces, a general familiarity signal exists that is shared across participants, stimuli, and mode of acquisition. The current study aims to explore the generalizability of this phenomenon by investigating the neural dynamics of cross-classification involving datasets for various stimulus types and sensory modalities. This allows for probing whether similar generalizable signals exist for recognition memory across different stimulus types and sensory modalities.

Replicating previous findings (Dalski et al., 2022a, 2022b; Li et al., 2022), neural signals for face stimuli generalized remarkably well starting from around 200 ms after stimulus onset. When training was performed on non-face datasets, an early (200-300 ms) and a late (post-400 ms) differentiation was observed over most regions of interest. Successful cross-classification for non-face (music and object/scene associations) datasets was more pronounced in the late period. Finally, a clear distinction was observed between familiar and remembered objects, with shared patterns only present for correctly remembered objects in the later time frame, while familiar objects also had shared patterns in the early period. Furthermore, classifiers trained on signals related to subjective familiarity and recollection (such as incorrect responses and false alarms) were effective in identifying genuine face-familiarity.

## Methods

### Test dataset

This analysis is based on the availability of openly accessible EEG datasets, which originated from different laboratories and were measured using different EEG devices and setups. In addition to the varying stimulus types, there were multiple other factors that varied across the experiments included in this study, such as the number of trials, repetition of unique stimuli, experimental task, and the uncertainty of the familiarity decisions (see also the **Further considerations** section of this paper). These varying and uncontrolled parameters can influence the signal-to-noise ratio and, as a consequence, the separability of the signals for the categories of interest. Previous research has shown that cross-classification performance is higher in the low-to-high signal-to-noise ratio direction (a phenomenon known as decoding direction asymmetry), and successful cross-classification supports the presence of shared information content in the signals (van den Hurk and op de Beeck, 2019).

Therefore, it was decided that event-related potentials (ERPs) from a single experiment, the Incidental Recognition study by Wiese et al. (Wiese et al., 2022), will serve as the “target” dataset for this analysis. There are several reasons for this choice. First, the sample size in this study was *n* = 22, which is one of the highest among the available studies. Second, the stimulus set in this study consisted of trial-unique images of highly personally familiar and unfamiliar faces, which were pre-experimentally familiar, salient, socially and emotionally relevant, and unambiguously categorizable as known or unknown. Third, the stimulus set was also participant-unique, which reduces the likelihood that signals for lower-level stimulus properties will contribute to the results of cross-participant classification analyses. Fourth, the images were presented for an extended period (1 second) that allowed for the testing of a longer time window. Finally, the trials of interest did not require any response from the participants, which means that neural patterns related to response preparation, execution, and monitoring should not contribute to the signal. The following section provides a brief overview of the experimental procedures.

#### Personally familiar and unfamiliar faces (Wiese et al., 2022)

The stimuli consisted of face photographs of unknown and highly personally familiar individuals (e.g., close friends, relatives). The 50-50 images were trial-unique, luminance-adjusted, and presented in grayscale for 1000 ms on a gray background. The familiar and unfamiliar identities were different for each participant. To keep participants engaged, a butterfly detection task was used, otherwise the experiment did not require any responses to the face stimuli. Trials were separated by a fixation cross, with a presentation duration of 1500-2500 ms. In addition to the trial-unique, familiar/unfamiliar stimuli, an additional familiar and an unfamiliar identity were presented, each with a single image repeated 50 times. These trials were not analyzed in the current study.

### Training datasets

#### Famous and unfamiliar (Wakeman and Henson, 2015)

This dataset contains data from 15 participants. The stimuli were greyscale photographs of famous (celebrity) and unfamiliar individuals of both sexes. The start of each trial was indicated by a fixation cross (400-600 ms), and the inter-stimulus interval was set to 1700 ms. The experiment included 300 famous and 300 unfamiliar trials, which were presented for an interval that varied between 800 and 1000 ms. In addition to famous and unfamiliar faces, scrambled face images were also presented during the experiment, but these trials were not included in the current analysis. Participants were required to make symmetry judgments for the images. For the purposes of this current study, familiar and unfamiliar trials were selected for each participant based on their responses in a post-experiment familiarity questionnaire.

#### An experimentally familiarized face and unknown faces (Sommer et al., 2021)

This dataset contains data from 15 participants. The experimental design followed the Jane/Joe paradigm described in Tanaka et al. (2006). Volunteers were familiarized with a target face (matched to their gender, “Joe” or “Jane”) for 10-60 seconds. The same male face was used for the “Joe” condition, and the same female face was used for the “Jane” condition, across all participants. The 10 unknown faces were also matched to the participant’s gender. Stimuli were presented for 500 ms, followed by a 500 ms blank period. Finally, a cue prompted the participant to indicate whether the target face was presented. A 500 ms presentation of a fixation cross separated the trials. Each face image was presented twice in 36 experimental blocks (for a total of 72 presentations of the familiarized face and 720 presentations of unfamiliar faces). The participants’ own face was also part of the stimulus set, but these trials were not included in the current analysis.

#### Familiar and unfamiliar music (Jagiello et al., 2019)

This dataset contains data from 10 participants. Volunteers passively listened to short, 750 ms long segments of familiar and unfamiliar (personally relevant and acoustically matched unknown) songs. The experiment was divided into 10 blocks, each with 100 familiar and 100 unfamiliar snippets, presented randomly, with an ISI between 1000 and 1500 ms. The participants were not required to make a response. Only averaged ERP data in the two conditions was openly available for this experiment.

#### Object-scene associations (Treder et al., 2021)

The dataset for this study contains data from 18 participants. The experiment consisted of eight runs, each of which was divided into pre-encoding delay, encoding, post-encoding delay, and retrieval blocks. During the delay blocks, participants were instructed to perform an even-odd decision task on numbers between 0 and 100. The encoding blocks consisted of 32 trials, each starting with a fixation cross presented for 1500 ± 100 ms. A unique, randomly selected object-scene combination was then displayed until the participant pressed a button (minimum 2500 ms, maximum 4000 ms). Participants were asked to make a plausibility decision for each combination (i.e., how likely it is for the combination to appear in real life). The retrieval blocks also consisted of 32 trials, starting with a fixation cross presented for 1500 ± 100 ms. In eight mini-blocks, either an object or a scene was presented as a cue (until the button was pressed, minimum 2500 ms, maximum 6000 ms). Participants were asked to indicate whether they remembered the corresponding associated pair from the encoding phase (e.g., a scene for an object cue, or an object for a scene cue). “Remember” responses were to be given if the memory was vivid enough to provide a detailed description of the associated item; this was followed up by an instruction to give a description of the target in 20% of the trials. For the purposes of this analysis, “remember” trials were relabeled as “familiar” and “forgot” trials were relabeled as “unfamiliar” to investigate the neural correlates of successful memory retrieval.

#### Object familiarity and recollection (Dimsdale-Zucker et al., 2022)

The dataset for this study contains data from 38 participants. The experiment included an encoding and a retrieval phase. During the encoding phase, 180 images of objects from the Bank of Standardized Stimuli (BOSS) were shown in color for 250 ms, each accompanied by an encoding question (“Would you find this item in a supermarket/convenience store?” or “Would this item fit in a fridge/bathtub?”). During the retrieval phase, one of the 180 previously presented images or 90 novel images was shown for 700 ms, followed by a “think” cue presented for 1700 ms. During this time, participants were instructed to withhold their responses in order to minimize movement-induced artifacts. Next, the object was shown again, and participants were asked to make a self-paced “remembered”, “familiar”, or “new” decision, as well as a confidence estimation on a scale from 1 (highly confident) to 4 (not at all confident). Finally, participants were asked to make a source memory judgement (fridge, supermarket, bathtub, convenience store). Trials were separated by a variable interstimulus interval of approximately 2 seconds. The presentation of stimuli and the order of the familiarity and source memory options were randomized for each participant. A 45–60-minute delay period separated the encoding and retrieval phases. For the purposes of this study, “old” items correctly identified as “familiar” or “remembered” were tested against correctly identified “new” items in two separate analyses. Additional trial selection and relabeling was performed to explore the neural signatures of false alarms and forgotten items, and to probe the similarity of neural signals for familiarity and recollection to those of familiar face processing.

For an overview of the datasets included in the analyses, see **Supplementary Information, Tables S1A** and **B**.

### Analyses

The analysis pipeline (see **Figure 1**) was based on procedures described in Dalski et al. (2022a). Within-experiment, leave-one-subject-out analyses were implemented on the test experiment to characterize the temporal evolution of the information content present for personally familiar and unfamiliar stimuli, and multivariate cross-classification analyses were conducted across experiments to probe the temporal dynamics of shared neural patterns related to memory recall, independent of stimulus type. Time-resolved MVPA, generalization across time, and sensor-space spatio-temporal searchlight analyses were performed.

**Figure 1.**
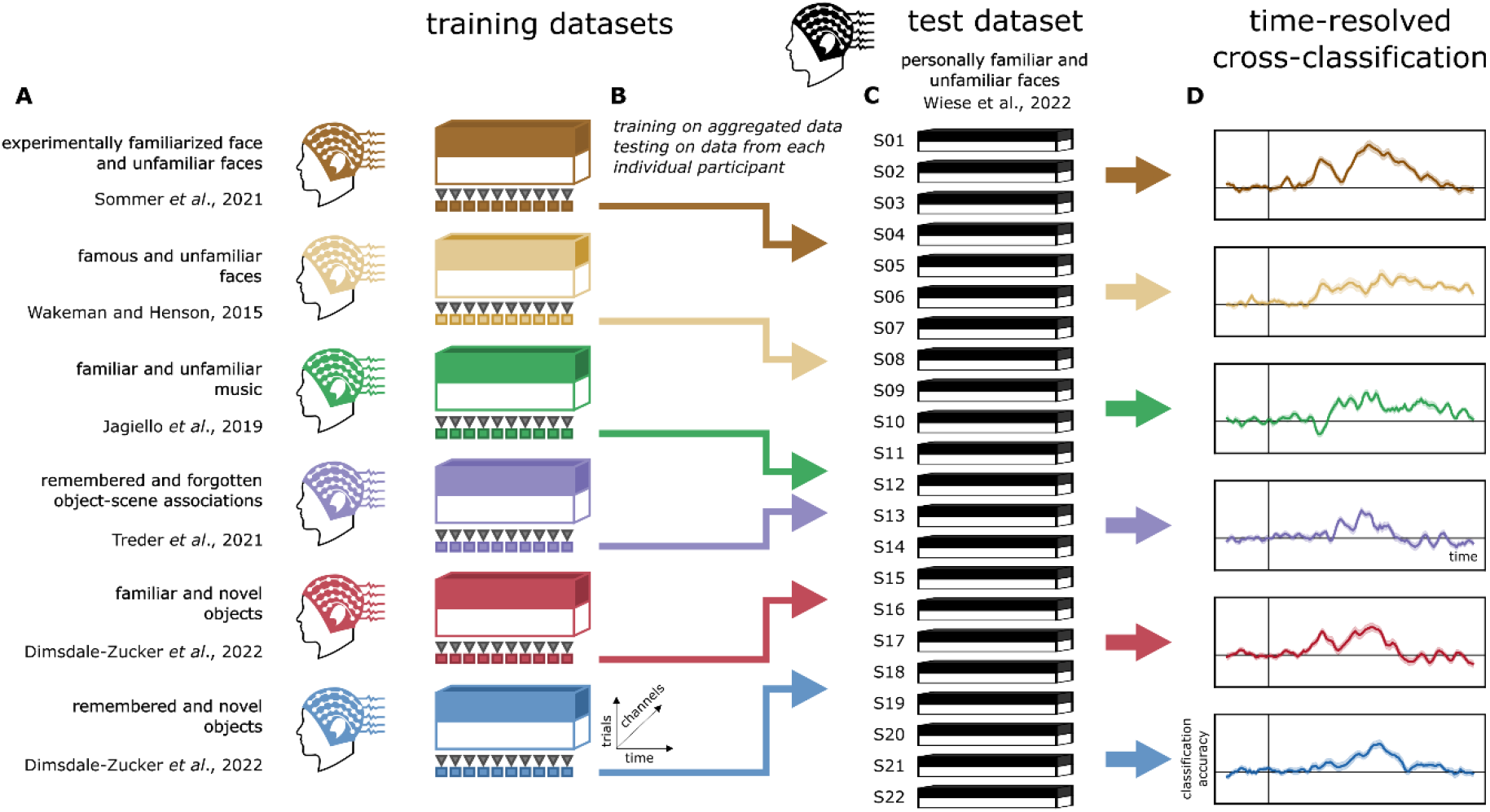
Analysis pipeline for time-resolved cross-classification. **(A)** ERP data (trials × times × channels) in each training dataset is aggregated across participants. **(B)** Linear discriminant analysis (LDA) classifiers are trained on data at each time point. **(C)** The fitted classifiers are then used to predict category membership in the test dataset, for each participant separately. **(D)** The classifier performance metrics are then aggregated to generate the sample-level cross-classification time course for each test dataset.

### Time-resolved multivariate pattern analysis

Time-resolved cross-classification was carried out on data form all channels, as well as pre-defined regions of interest. The ROIs were based on Ambrus et al. (2021, 2019b) and Dalski et al. (2022a). Six scalp locations along the medial (left and right) and coronal (anterior, center, and posterior) planes were defined and used for training and testing in separate analyses. Cross-classification was performed on common channels in the training and test dataset (see **Supplementary Information, Figure S1**). For each test dataset, linear discriminant analysis (LDA) classifiers were trained on aggregated data from all participants at each time point. These fitted classifiers were then used to predict stimulus familiarity in the test dataset for each trial in each participant.

### Spatio-temporal searchlight

All sensors were systematically tested separately by training and testing on data originating from the given channel and adjacent electrodes. For each channel and its neighbors, a time-resolved analysis was conducted.

### Generalization across time

Temporal generalization analysis was used to investigate the temporal organization of information-processing stages. For this, the classifiers were trained on data from each time point in the test experiments; these classifiers were then tested on the data at every timepoint in the test experiment. The results were organized in a cross-temporal (training-times × testing-times) classification accuracy matrix. Similar information-processing may be indicated where classifiers successfully generalize from one timepoint to another across the experiments (King and Dehaene, 2014).

### Statistical testing

The analyses were conducted on common-average-referenced, baseline-corrected (-200 to 0 ms), and down-sampled (100 Hz) data. Only trials with correct responses were included in the main analyses (where applicable). Trial counts were balanced at both training and testing by under-sampling to the minimum trial count in the classes of interest, in each participant. A moving average of 30 ms (3 consecutive time points) was applied to all decoding accuracy data at the participant level (Ambrus et al., 2019b; Dalski et al., 2022a). For ROI-based time-resolved analyses, classification accuracies were entered into two-tailed, one-sample cluster permutation tests (10,000 iterations) against chance (50%). For the temporal generalization and searchlight analyses, two-tailed spatio-temporal cluster permutation tests were used against chance level (50%), with 10,000 iterations.

Statistical analyses were conducted using python, MNE-Python (Gramfort et al., 2014), scikit-learn (Pedregosa et al., 2011) and SciPy (Virtanen et al., 2020).

## Results

### Characterization of the test dataset

To characterize the evolution of the information content in the test dataset, within-experiment, leave-one-subject-out (LOSO) classification analyses were conducted on the personally familiar and unfamiliar faces experiment (Wiese et al., 2022). The time-resolved classification analysis revealed strong, sustained clusters beginning around 200 ms in all regions of interest (cluster *p* < 0.0001, **Figure 2A**, see **Supplementary Table 1A**). Likewise, the searchlight analysis yielded a single cluster encompassing all sensors (cluster *p* < 0.0001, **Supplementary Table 5A**). Temporal generalization analyses also showed sustained effects starting around 200 ms (cluster *p*-values < 0.0001, Figure 2B, see **Supplementary Table 3A**).

**Figure 2.**
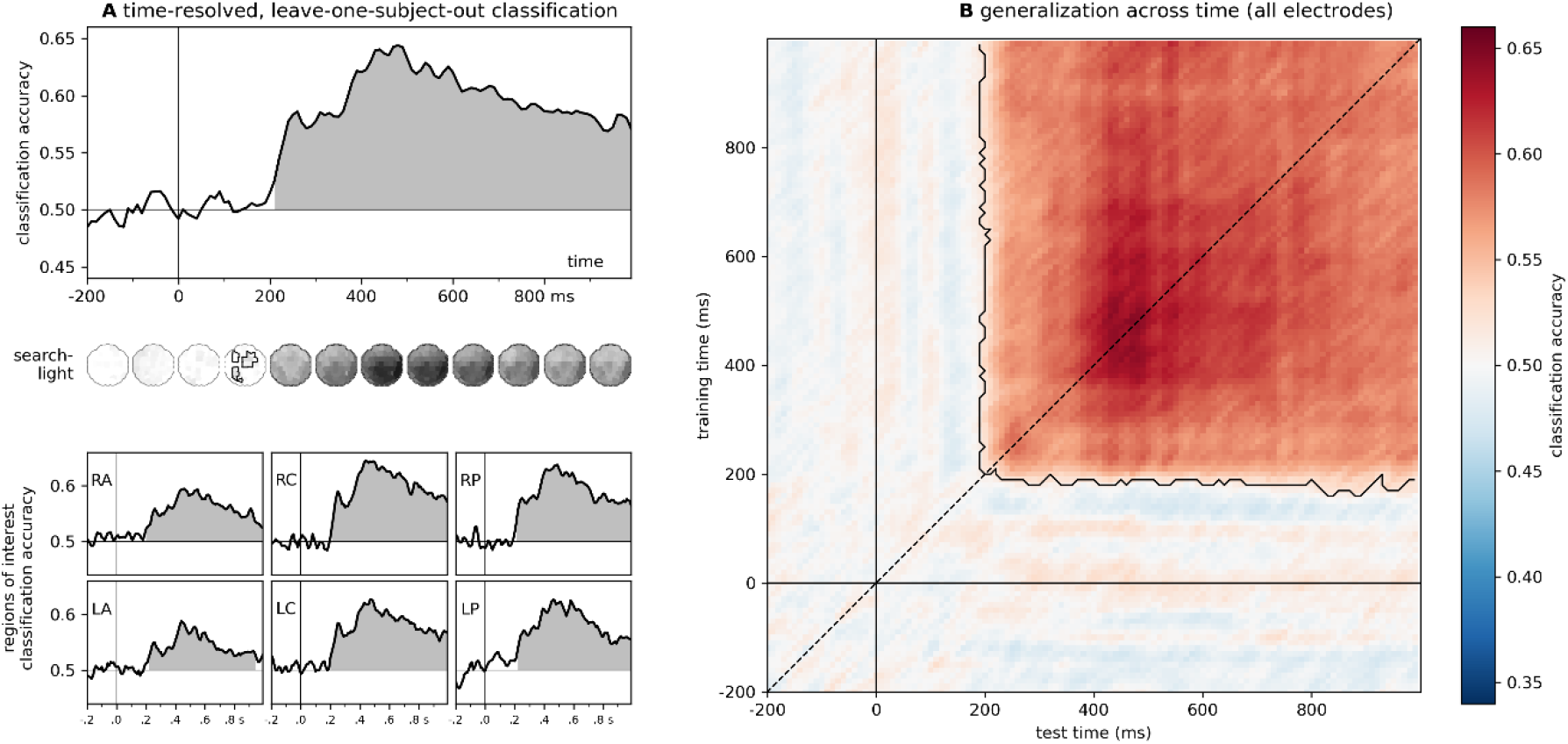
Familiarity classification in the test dataset. Within-experiment, leave-one-subject-out classification for the personally familiar and unfamiliar faces (Wiese *et al*. 2022) experiment. A) *Time-resolved classification* over all electrodes and pre-defined regions of interest. Shaded regions denote significant clusters (two-sided cluster permutation tests, *p* < 0.05). For detailed statistics, see **Supplementary Table 1A**. Spatio-temporal searchlight results are shown as scalp maps, with classification accuracy scores averaged in 100 ms steps. Regions forming part of positive clusters (*p* < 0.05) are marked. Statistics calculated using spatio-temporal cluster permutation tests. B) *Generalization across time*. Two-sided cluster permutation test (p < 0.05). RA/LA: right/left anterior, RC/LC: right/left central, RP/LP: right/left posterior

### Cross-experiment classification

Cross-experiment time-resolved classification over all electrodes yielded significant effects for all six datasets (**Figure 3, Supplementary Information Figure S2**). Sustained, robust effects were observed from ca. 200 ms for both face-familiarity datasets (experimentally familiarized faces, **Figure 3**A: 200-340 ms, cluster *p* = 0.013, 370-800 ms, cluster *p* < 0.0001; famous faces, **Figure 3**B: 220 - 990 ms, cluster *p* = 0.0002), while for other stimulus types, successful cross-classification was not indicated before 300 ms post-stimulus onset (familiar music, **Figure 3**C: 300-890 ms, cluster *p* = 0.0001, 920-970 ms, cluster *p* = 0.035; object-scene associations, **Figure 3**D: 330-630 ms, cluster *p* < 0.0001; remembered objects, **Figure 3**F: 410-630 ms, cluster *p* < 0.0001). Interestingly, the exception to this was the case of familiar objects (**Figure 3**E), where the start of the significant cluster (200-620 ms, cluster *p* < 0.0001) coincided with that of the leave-one-subject-out classification in the test dataset (see also **Supplementary Information 1**).

**Figure 3.**
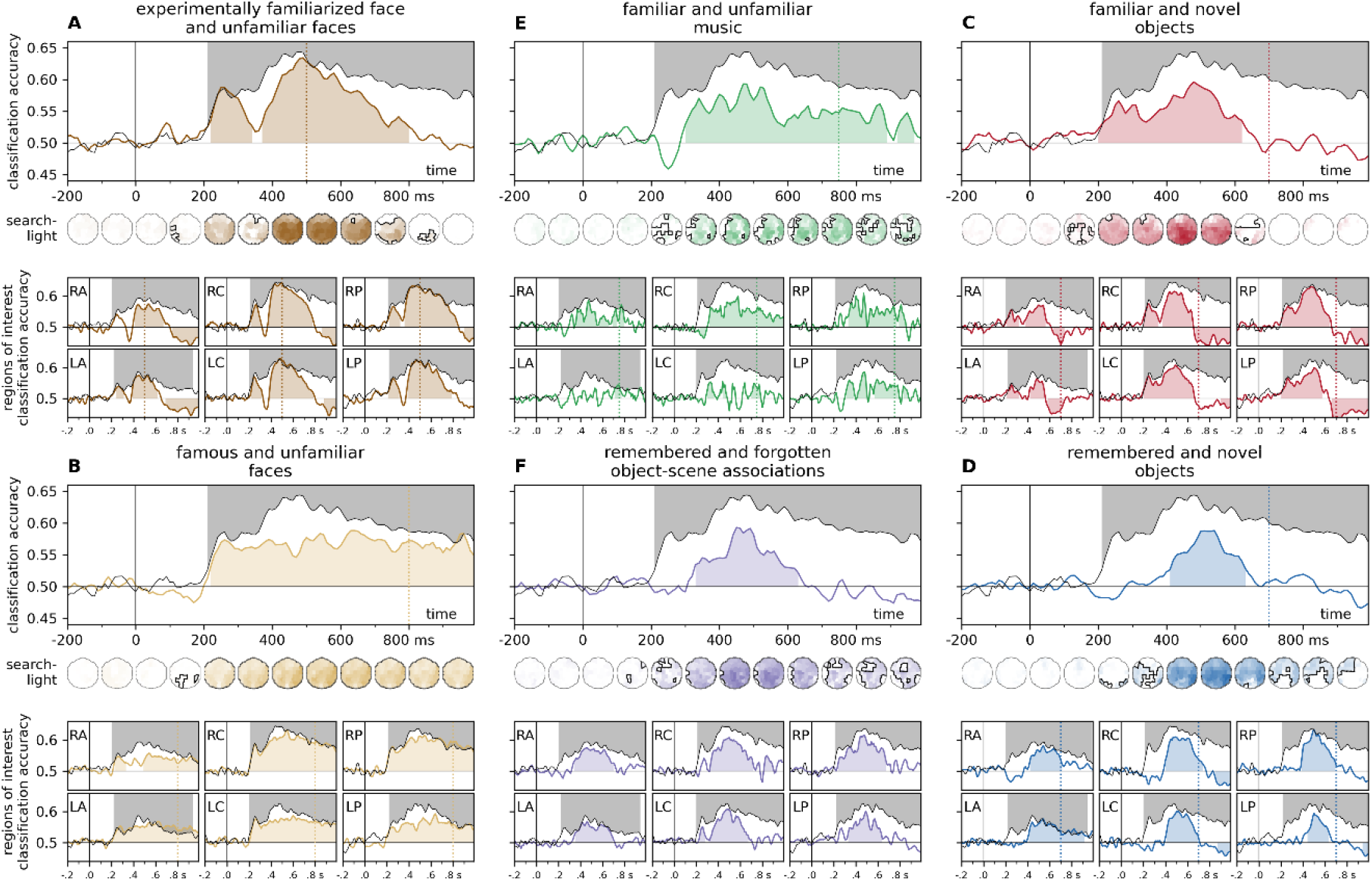
Time-resolved cross-experiment classification accuracies over all electrodes and pre-defined regions of interest. Classifier training was performed on EEG data from a range on memory tasks, and classification accuracy was tested on ERPs for highly personally familiar and unfamiliar faces (Wiese *et al*.). For comparison, the results of the leave-one-subject-out classification analysis for this test dataset are overlayed in black. **Left. Face memory**. Classifiers trained on ERPs for an experimentally familiarized face and a set of unfamiliar faces (Sommer *et al*., **A**, dark brown) and faces of pre-experimentally familiar celebrities and unknown faces (Wakeman and Henson., **B**, light brown) **Middle. Non-face memory**. Classifiers trained on ERPs for short segments of pre-experimentally familiar and unfamiliar music (Jagiello *et al*., **C**, green) and experimentally learned object/scene associations (Treder *et al*., **D**, purple). Significant clusters emerged most prominently in the late, post-300 ms time window. For music stimuli, the most consistent effect was observed over the right central ROI, while for scene-object associations, frontal and central regions also yielded sustained significant clusters. **Right. Objects, familiar and remembered**. Classifiers trained on ERPs for images of objects (Dimsdale-Zucker *et al*.) participants indicated as familiar (**E**, red) or remembered (**F**, blue). Significant clusters for both conditions included the late, ca. 400-600 ms window, while the early, ca. 200-300 ms window was only flagged in the ‘familiar’ condition. Statistics were calculated using two-sided cluster permutation tests (*p* < 0.05), shaded regions denote significant clusters. The end of the stimulus presentation window is marked by vertical dotted lines. For detailed statistics, see **Supplementary Table 1**. Spatio-temporal searchlight results are shown as scalp maps, with classification accuracy scores averaged in 100 ms steps. Regions forming part of a positive clusters are marked. Statistics were calculated using spatio-temporal cluster permutation tests. For detailed results, see **Supplementary Table 5**. RA/LA: right/left anterior, RC/LC: right/left central, RP/LP: right/left posterior

The results of the spatio-temporal searchlight analyses yielded sustained effects that encompassed the majority of the sensors, peaking between 460 and 560 ms. Training on experimentally familiarized faces yielded a positive (190-850 ms, cluster *p* < 0.0001) and a negative (690-990 ms, cluster *p* = 0.023) cluster, while a single, sustained (190-990 ms, cluster *p* < 0.0001) cluster was seen in the case of famous faces. Music stimuli and object-scene associations both yielded sustained positive clusters (music: 270-990 ms, cluster *p* < 0.0001, object-scene associations: 200-990 ms, cluster *p* < 0.0001). A robust early positive cluster emerged for familiar objects (190-660 ms, cluster *p* < 0.0001) which was followed by a later negative cluster (560-990 ms, cluster *p* = 0.0046). For remembered objects, the cluster permutation test flagged a large time interval (220-990 ms, cluster *p* < 0.0001) as significant, although only a handful of posterior sensors were part of the cluster between 200 and 400 ms.

Region-of-interest-analyses have shown that for most stimulus types, the most prominent and consistent effects were observed over the right central and posterior ROIs (**Figure 3**, see also **Figure 4A and 4B**). Negative classification accuracies (i.e., cases of significant, below-chance classification) were seen in some training datasets, these were mostly restricted to the post-500 ms interval. For detailed statistics, see **Supplementary Table 1**.

**Figure 4.**
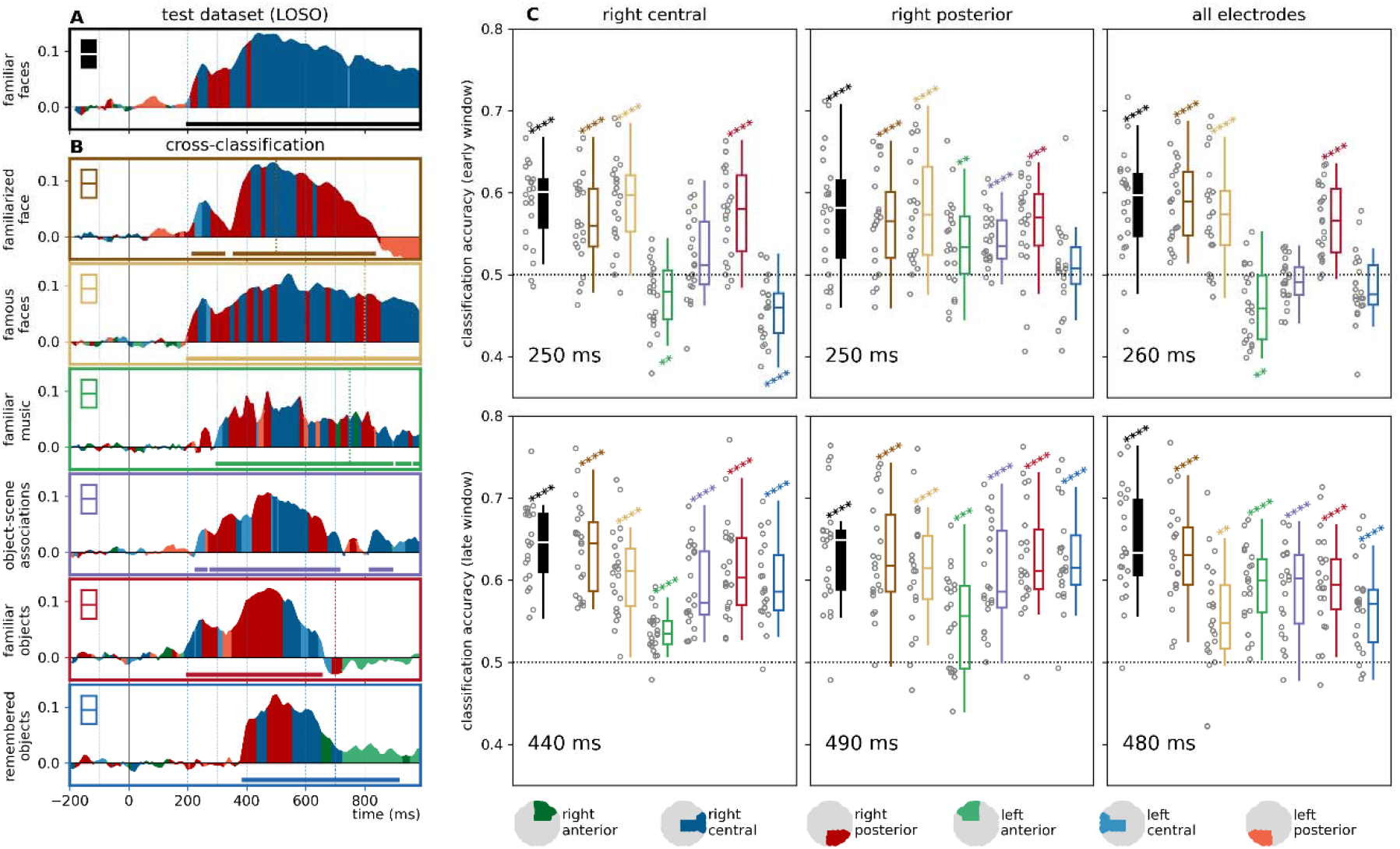
Cross-classification accuracy in early and late phases. Left panel: classification accuracy at each time-point, at the region of interest of highest classification accuracy, compared to mean baseline classifier performance, **(A)** in the leave-one-subject-out analysis for the test dataset, and **(B)** the cross-classification analyses. Horizontal significance markers indicate if the time-ROI datapoint belonged to a significant cluster based on the results of the cluster permutation test reported in **Figures 2** and **3**. Highest decoding accuracies were seen in the right central and right posterior regions of interest. **(C)** Classification accuracies in the time-resolved cross-classification analyses in early and late phases. The time points depicted correspond to accuracy peaks in the results of the LOSO analysis for the test dataset. Results in the right central and posterior ROIs, as well as all electrodes, are depicted here. Results of all ROIs can be found in **Supplementary Information S3**. Statistics: two-sided one-sample t-tests, *p*_uncorrected_ < 0.05*, < 0.01**, < 0.001***, < 0.0001**** Boxplots: first, second (median), and third quartiles; whiskers with Q1 − 1.5 IQR and Q3 + 1.5 IQR, with horizontally jittered individual classification accuracies.

Region-of-interest analyses were followed up by examining cross-classification accuracies for the different training datasets at time-points with peak within-experiment leave-one-subject-out decoding accuracies in the test dataset. In the early window, significant positive effects were seen only in the case of the face-familiarity datasets, and in the case of objects correctly identified as “familiar”. In contrast, classification accuracies at the late peak consistently yielded significant effect in all cross-classification analyses (**Figure 4C, Supplementary Figure 3A**).

Temporal generalization analyses over all electrodes generally yielded rectangular cross-classification matrices (**Figure 5**). In the case of experimentally familiar faces an early positive and a later negative cluster emerged (train interval: 90 to 830 ms, test interval: 190 to 990 ms, cluster *p* < 0.0001; train interval: 820 to 1290 ms; test interval: 220 to 990 ms, cluster *p* = 0.014). For famous faces, a single sustained cluster was observed (train interval: 200 to 1290 ms, test interval: 200 to 990 ms, cluster *p* = 0.0002). Music stimuli yielded a large positive cluster (train interval: 290 to 1100 ms, test interval: 200 to 990 ms, cluster *p* < 0.0001) in addition to later positive (train interval: 1130 to 1490 ms, test interval 190 to 990 ms, cluster *p* = 0.0029) and a number of smaller, weak effects that also included periods in the baseline. In the case of object-scene associations, two early positive clusters (train interval: -20 to 210 ms, test interval: 300 to 990 ms, cluster *p* = 0.0056; train interval: 310 to 640 ms, test interval: 230 to 990 ms, cluster *p* = 0.0005), a weaker negative cluster (train interval: 650 to 1030 ms, test interval: 200 to 710 ms, cluster *p* = 0.0425) and a late, weaker positive cluster (train interval: 1130 to 1290 ms, test interval: 290 to 780 ms, cluster *p* = 0.0249), were seen. Both familiar and remembered objects yielded an earlier positive and a later negative cluster (familiar objects: train interval: 130 to 770 ms, test interval: 160 to 990 ms, cluster *p* < 0.0001; train interval: 850 to 1290 ms, test interval: 240 to 990 ms, cluster *p* = 0.0014; remembered objects: train interval: 400 to 760 ms, test interval: 280 to 990 ms, cluster *p* = 0.0046; train interval: 760 to 1290 ms, test interval: 180 to 990 ms, cluster *p* = 0.0002). For the results and the detailed statistics of the temporal generalization analyses in the regions of interest, see **Supplementary Table 3**.

**Figure 5.**
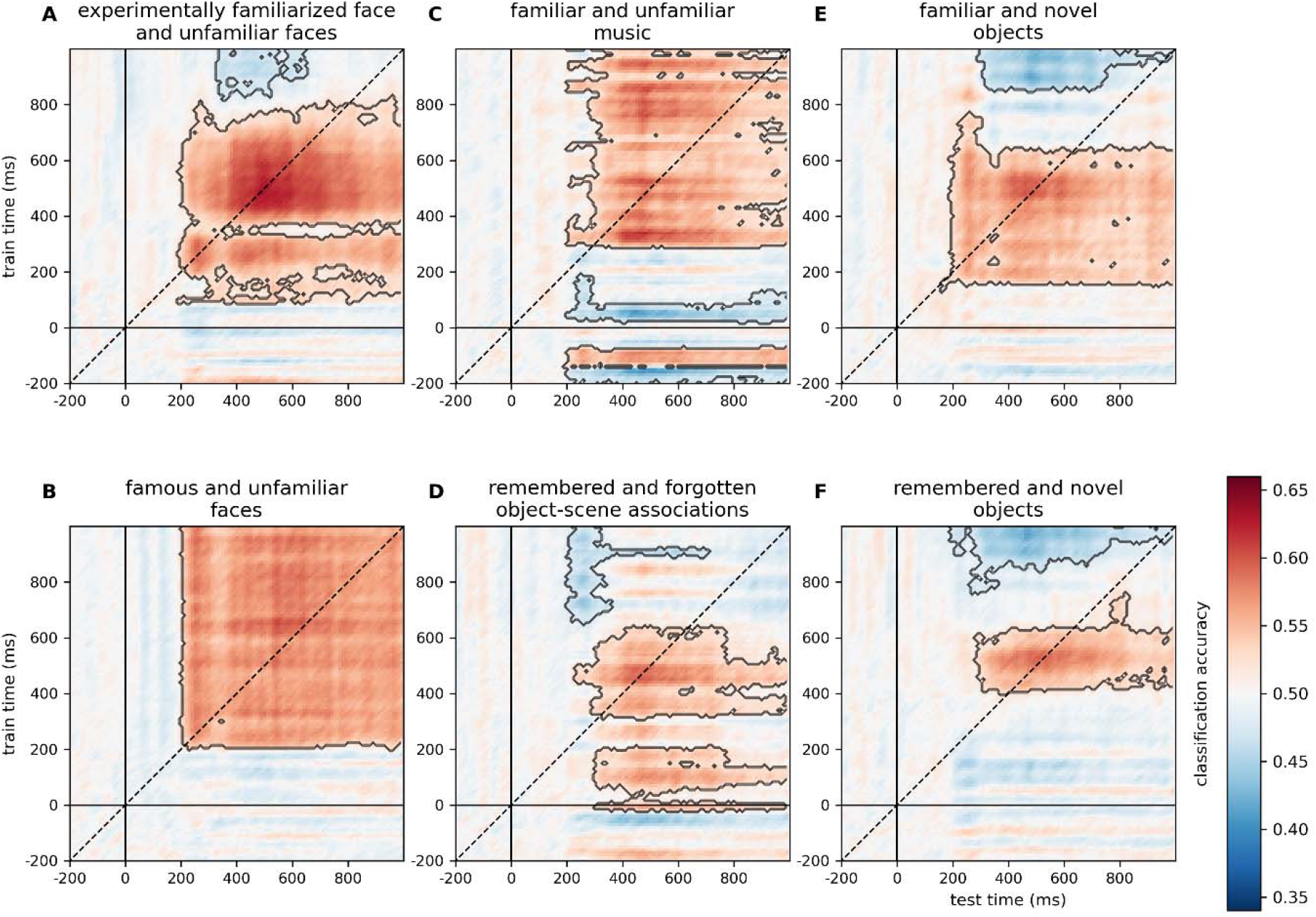
Cross-experiment temporal generalization analysis over all electrodes. Two-sided cluster spatio-temporal permutation test (p<0.05). For the results of the temporal generalization analyses separately in the different ROIs, see **Supplementary Table 3**.

### Subjective effects in familiarity and recollection

The object familiarity/recollection dataset was subjected to further analyses. To test the cross-experiment representational dynamics of false alarms and forgotten items, these were relabeled as “familiar”, while keeping the “unfamiliar” labels of correctly rejected novel items. For forgotten items, old items incorrectly categorized as new were relabeled as “familiar”, while the “unfamiliar” label for novel items were kept (see **Supplementary Information, Table S1B**). Classifiers were then trained on these newly constructed datasets and tested on the ERPs for personally familiar and unfamiliar faces.

Time-resolved cross-classification over all electrodes in the case of false alarms for familiar objects yielded two positive clusters (190 – 340 ms, cluster *p* = 0.0006; 360 – 590 ms, cluster *p* < 0.0001). For false-alarm-remembered objects, three positive clusters were observed (280 - 330 ms, cluster *p* = 0.0308; 370 - 460 ms, cluster *p* = 0.0155, and 480 - 800 ms, cluster *p* < 0.0001). Spatio-temporal searchlight analysis yielded one positive cluster (180 - 940 ms, cluster *p* < 0.0001) (**Figure 6A**). A single late positive cluster was observed in the case of forgotten vs. novel objects (370 - 560 ms, cluster *p* = 0.0056) in the time-resolved cross-classification analysis, while the spatio-temporal searchlight yielded a cluster (190 - 990 ms, cluster *p* < 0.0001) that also encompassed earlier time points (**Figure 6B**).

**Figure 6.**
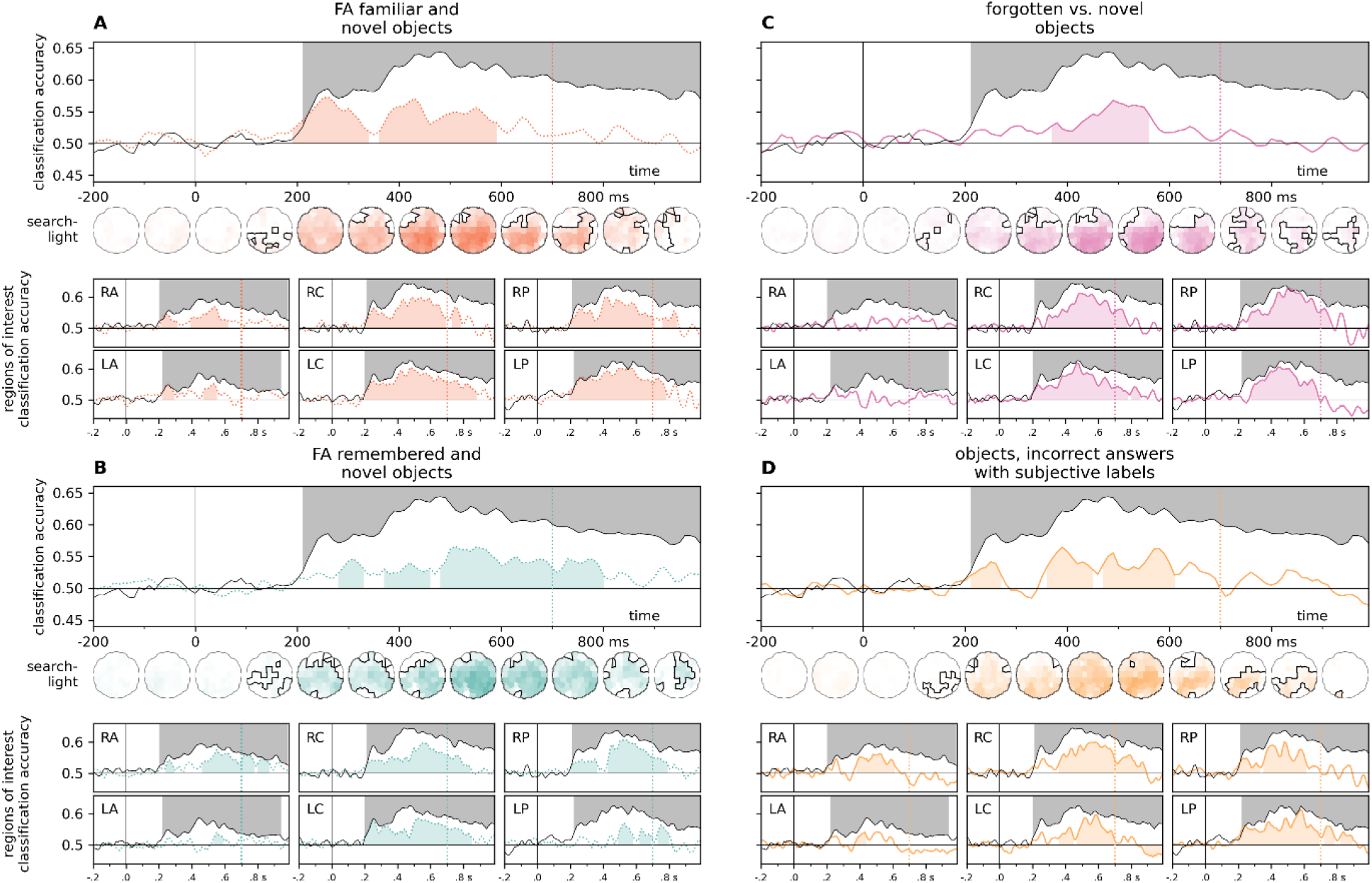
Time-resolved cross classification for false alarms, forgotten items, and subjective old and new, but incorrect answers in the object recognition dataset. These analyses required the relabeling of specific trials in the training dataset **(A) False alarms for familiar objects**. familiar: new item, response: familiar. unfamiliar: new item, response: new. **(B) False alarms for remembered objects**. familiar: new item, response: remembered. unfamiliar: new item, response: new **(C) Forgotten objects. familiar:** old item, response: new. unfamiliar: new item, response: new. **(D) Subjective old/new answers in trials with incorrect responses**. familiar: new item, response: old. unfamiliar: old item, response: new. Results of the leave-one-subject-out classification for personally familiar and unfamiliar faces is presented in black for comparison (two-sided cluster permutation tests, *p* < 0.05). Spatio-temporal searchlight results are shown as scalp maps, with classification accuracy scores averaged in 100 ms steps. Sensors and time points forming part of significant clusters are marked (Two-sided spatio-temporal cluster permutation tests, *p* < 0.05). RA/LA: right/left anterior, RC/LC: right/left central, RP/LP: right/left posterior

Time-resolved analysis for forgotten objects over all sensors yielded a late positive cluster (370 - 560 ms, cluster *p* = 0.0056), while a positive cluster (160 - 990 ms, cluster *p* < 0.0001) was observed in the results of the searchlight analysis. Interestingly, both ROI-based and searchlight results have shown weaker effects over frontal electrodes (**Figure 6C**).

To test the neural signals for subjective (but incorrect) recall, only trials with erroneous responses were retained. ERPs for these trials were then labeled according to the participants’ response (i.e., incorrect “old” trials were labeled “familiar”, and incorrect “new” trials were labeled “unfamiliar”). Remarkably, the time-resolved analysis also yielded positive clusters at both early and late time points (210 - 270 ms, cluster *p* = 0.0262; 360 - 450 ms, cluster *p* = 0.006; 470 - 610 ms, cluster *p* = 0.004). The searchlight analysis also yielded a positive cluster with an early onset (190 - 920 ms, cluster *p* < 0.0001), as well as a weak late negative cluster (840 - 990 ms, cluster *p* = 0.0416).

To explore the similarity of neural signals for familiarity with, and recollection of objects to those of familiar face processing, classifiers were trained to categorize ERPs for objects judged as familiar, and objects judges as remembered. These classifiers were then tested on ERPs elicited by familiar faces. Time-resolved cross-classification (**Figure 7**) revealed an early (190-410 ms, cluster *p* = 0.002) cluster in favor of familiarity, while a late cluster favored recollection (660 - 840 ms, cluster *p* = 0.003). Similarly, the spatio-temporal searchlight analysis yielded early evidence for familiarity (190 - 530 ms, cluster *p* = 0.002), and late evidence for recollection (370 - 990 ms, cluster *p* < 0.0001). For detailed statistics, see **Supplementary Table 2** and **Supplementary Table 6**.

**Figure 7.**
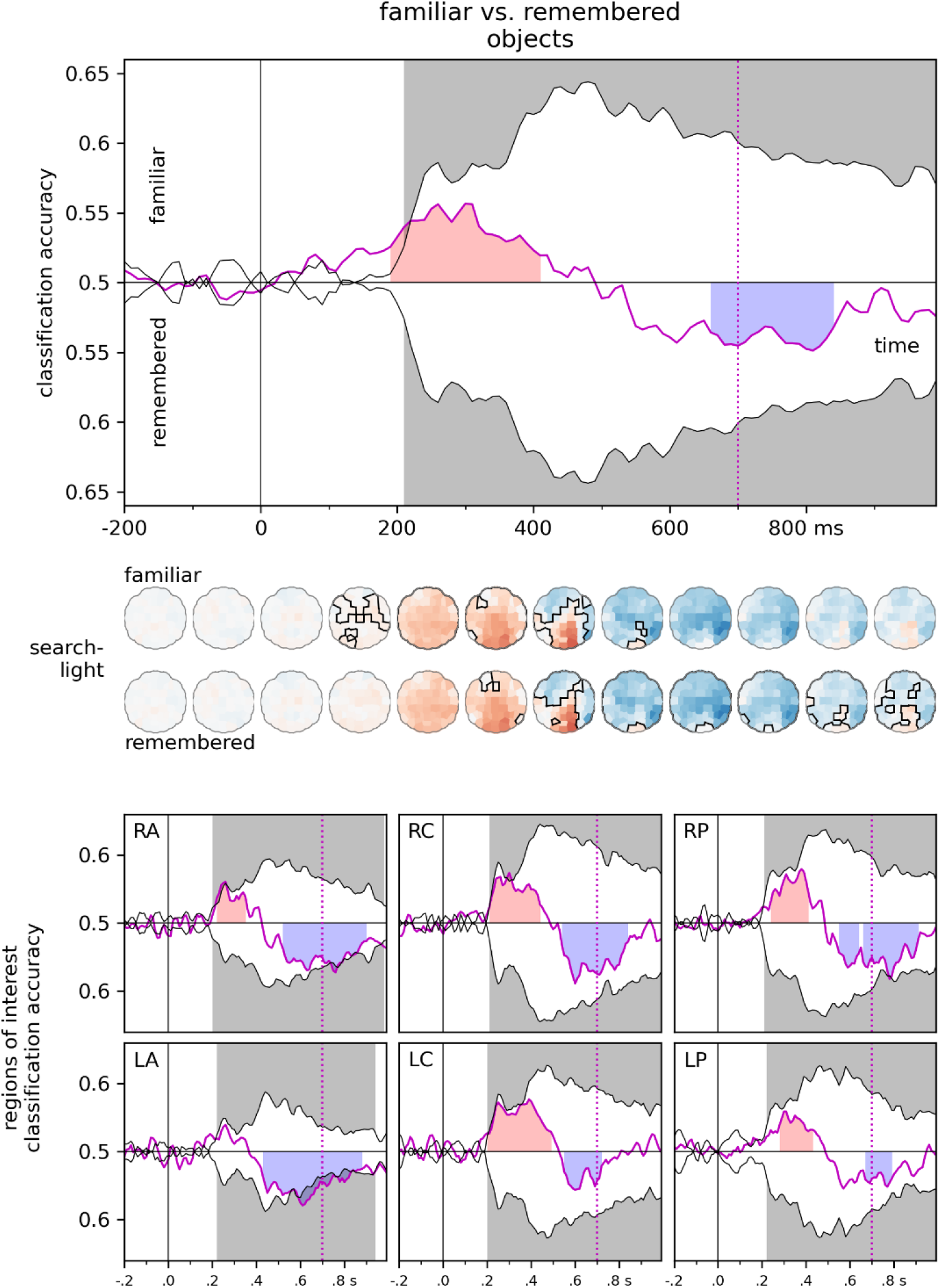
Familiar vs. Remembered objects. Classifiers were trained to categorize ERPs for familiar and remembered objects and were tested on ERPs for familiar faces. Regions shaded in red denote significant evidence for familiarity, regions shaded in blue denote significant evidence for recollection. Results of the leave-one-subject-out classification for familiar faces is presented in black for comparison. (Two-sided cluster permutation tests, *p* < 0.05). Spatio-temporal searchlight results are shown as scalp maps, with classification accuracy scores averaged in 100 ms steps. Sensors and time points forming part of the significant cluster evidencing familiarity are shown in the top row, sensors and time points forming part of the significant cluster evidencing recollection are shown in the bottom row. (Two-sided spatio-temporal cluster permutation tests, *p* < 0.05). RA/LA: right/left anterior, RC/LC: right/left central, RP/LP: right/left posterior

## Discussion

The aim of this study was to probe the shared signals of memory functions using a data-driven approach. Much of the previous work in this area focused on canonical ERP components in a priori spatial and temporal windows. While this has led to the identification of a number of components related to memory processes, it is also limited in a number of ways. For example, the timing of certain components may shift depending on experimental factors, relevant signals might be mixed with those of other cognitive processes, and signatures of memory processes that do not confirm to canonical configurations might be overlooked.

Here, time-resolved cross-experiment classification of ERP signals (Dalski et al., 2022a, 2022c, 2022b; Li et al., 2022) was used to find shared information content indicative of similar underlying memory-related processes across participants, stimulus modalities and tasks, and to probe how these signals unfold over time. Classifier training was conducted on data from experiments investigating face familiarity (Sommer et al., 2021; Wakeman and Henson, 2015), object familiarity and recollection (Dimsdale-Zucker et al., 2022), familiar and unfamiliar music (Jagiello et al., 2019), and remembered and forgotten object-scene associations (Treder et al., 2021). ERPs for highly familiar and unfamiliar faces comprised the test dataset (Wiese et al., 2022).

The main findings of these analyses were the following. 1) Cross-classification performed on all training datasets yielded sustained significant effects; 2) depending on the training dataset, dissociations between an early (ca. 200 to 400 ms) and a late (ca. 400 to 600 ms) time window were observed; 3) familiar and remembered objects differed substantially in the early phase; 4) subjective recognition judgements also yielded positive patterns; and 5) significant late negative decoding was observed for some training datasets.

### Early and late effects

Broadly defined early and late effects, such as the FN400 and the LPC, have long been investigated in memory research. Person-perception research uses a different nomenclature, somewhat independent from that of the rest of the field, such as the N170, the N250, and the SFE (but see for example Herzmann and Sommer, 2010; MacKenzie and Donaldson, 2007; Nessler et al., 2005). The fact that face-specific components, such as the N250, have also been described for familiar objects trained for expertise, as well as for personally familiar and newly learned objects (Pierce et al., 2011; Scott et al., 2008, 2006) further complicates the picture. Due to the considerable differences in the timing and topographical distribution across studies using different stimulus types and experimental paradigms (Dimsdale-Zucker et al., 2022; Kwon et al., 2022), the relationship between these effects remains unclear.

Another factor to consider is the functional separation between components described for different time windows. While an extensive body of literature exists evidencing the differential modulation of the NF400 and the LPC effects, such evidence for early and late face-related effects, that share a similar topographic distribution, has started to emerge only recently. Dalski et al. (2022a) reported sustained cross-dataset temporal generalizability of face-familiarity information between ca. 200 and 600 ms following stimulus onset, a finding that was interpreted as an indication of a single automatic processing cascade that is maintained over time. In this present analysis, temporal cross-classification on face-datasets (**Figure 5**A) yielded very similar results. Nevertheless, while similar in scalp distribution, evidence exists that suggests that the early (200 - 400 ms) and late (>400 ms) effects might be functionally distinct. Wiese and colleagues found that ERP markers of familiarity are modulated differentially in these two time-windows: while the N250 (200 – 400 ms) is not substantially affected by attentional load, the sustained familiarity effect (SFE, >400 ms) relies on the availability of attentional resources (Wiese et al., 2019a). Cross-classification evidence comes from Li et al. (2022) who found that cross-classification accuracy for famous faces correlated with the level of familiarity in the ca. 400 – 600 ms window only, and Dalski et al. (2022c), who reported differential modulation in the early and late windows as a function of the task.

Although the N250 is commonly interpreted as a face-specific ERP component that reflects access to the visual representation of a face, it has been suggested that it also indexes enhanced access to post-perceptual representations. The SFE is theorized to reflect the integration of visual information with person-related (for example, semantic and affective) knowledge (Wiese et al., 2019b), or an additional elaboration phase in the perceptual face recognition process (Wiese et al., 2019a).

The results of this present analysis, when training on non-face datasets, also argue for at least a partial functional separation between these two phases. In the right occipito-temporal regions of interest, where within-experiment classification yielded the highest decoding accuracies, a marked dissociation between early and late effects can be observed. Early effects (**Figure 4**C, top row) were unambiguously present in the case of face training datasets, and were not significantly above chance in other comparisons, with the important exception of familiar vs. novel objects (see the **Familiarity and Recollection** section below). In contrast, significant late effects were observed in all comparisons (**Figure 4**C, bottom row).

### Stimulus material

In all comparisons, strong effects were observed in the approximately 300-400 to 600 ms time window, and these effects encompassed a large number of channels and regions of interest (see **Supplementary Information Figure S3B**).

#### Faces

In Dalski, Kovács, & Ambrus (Dalski et al., 2022a) we reported on the generalizability of face-familiarity signals across experiments, where familiarization was achieved either perceptually, via media exposure, or by personal interaction (Ambrus et al., 2021). The significant cross-experiment face-familiarity classification involved all three datasets, predominantly over posterior and central regions of the right hemisphere in the 270–630 ms time window. This current study replicates this finding on three independent datasets, providing further evidence for the existence of a robust, general face-familiarity signal.

#### Familiar and unfamiliar music

The authors of the report (Jagiello et al., 2019) noted that the EEG responses they observed were similar to those commonly observed in old/new memory retrieval paradigms, suggesting that similar mechanisms may play a role in their generation. This present analysis supports this observation. Evidence for sustained positive cross-classification emerged ca. 300 ms after stimulus onset, most prominently over right central and posterior sites.

The interpretation of these findings needs to take into account that the music material in this experiment was tied also tied to person-knowledge in at least two different ways. First, the familiar music presented was chosen to be personally relevant, as such, effects of personal semantics and autobiographical access cannot be ruled out (Jakubowski and Ghosh, 2021). Second, the stimulus set comprised of contemporary popular music. That means, the auditory stimuli might be also associated with access to biographic knowledge of the artists, as well as visual imagery from the accompanying music videos. As evidence exists for the cross-domain activation of related representations, priming being one example (Ambrus et al., 2019a; Koelsch et al., 2004), our understanding would benefit from systematically investigating how these factors relate to shared neural patterns of memory functions.

Although large and sustained effects were observed, in contrast to pre-experimentally familiar faces and experimentally familiarized objects, significant cross-classification effects did not manifest earlier than ca. 300 ms after stimulus onset. Two mutually non-exclusive hypotheses present themselves. Music is inherently sequential in nature, as such, in contrast to visual information, the generation of a recognition signal might be delayed by the time needed for the accumulation of the necessary evidence. Studies suggest that a 500 ms long excerpt is needed to reliably identify familiar and unfamiliar music (Filipic et al., 2010; Tillmann et al., 2014); in a closed set, such as the dataset used here, 100 ms has been shown to be sufficient (Schellenberg et al., 1999). Indeed, the authors measured pupillary responses concurrently with EEG and found an early familiarity effect starting around ca. 100 ms, for which they did not observe a corresponding early ERP component. The temporal generalization matrix in this present analysis, however, yielded significant clusters in central and posterior ROIs for training times between ca. 100 and 170 ms. Although caution must be exercised in taking this weak effect at face value, for the sake of completeness, it is nevertheless reported in **Supplementary Information 4**. It should be noted that while both the train and test datasets used long-term, personally relevant stimuli, they differed in the type of stimulation they employed. The test dataset used visual stimuli, while the train dataset used auditory stimuli. The delay in cross-decodable early effects may also suggest that early signals of recognition may vary depending on the type of stimulation.

#### Object-scene associations

Successful, sustained effects were observed approximately 330 ms following stimulus onset over all electrodes, while significant effects over central and posterior clusters were seen as early as 230 ms. Frontal effects first emerged around 350 ms following stimulus onset. It is worth noting that, in contrast to the other experiments analyzed in this study, all stimuli in the test phase were presented in the learning phase and the “familiar” and “unfamiliar” labels were based on whether the participant reported remembering them or not. This means that there were two ways to generate “forgotten” responses: either the participant failed to recall both the cue and the target stimuli, or they remembered (or felt familiarity with) the cue but failed to recall the paired associate. In contrast, “remembered” responses, indicating successful recall, required the participants to remember both the cue and the related target stimulus. How much the recognition of the cue and the successful recall of the target contributed to successful cross-classification is difficult to disentangle, but for reasons detailed in the following sections, it is possible that early peaks over posterior and central regions of interest are more related to a sense of familiarity with the cue, while the later effects are more related to the recollection of the paired associate.

#### Experimentally familiarized objects

Training on ERPs for familiar objects and testing on personally familiar and unfamiliar faces yielded strong, early (200 ms) effects. In fact, beside the two face-ERP datasets, this condition yielded the most stable early patterns across all regions of interest. As the stimulus material in these two studies was vastly different, the effect of low-level image properties can be ruled out with confidence. This may imply that by this time point, for both faces and objects, the visual information is processed in adequate detail to elicit a stimulus-independent familiarity signal. In stark contrast, cross-classification patterns for images of objects judged as remembered first emerged around 400 ms post-stimulus onset. A more detailed discussion is given in the following section.

### Familiarity and recollection

As hinted above, while both early and late cross-classification accuracies in the case of familiarity for newly learned objects and personally familiar faces were remarkably high, no early effect was seen in recollection for objects. It is important to emphasize that this lack of early cross-classification between remembered objects and familiar faces does not necessarily indicate the absence of recognition information present in this early period, only that there were no utilizable shared neural signals between the processes by which recognition is mediated for remembered objects and familiar faces in this early stage. Late signals, on the other hand, were highly informative both in the case of familiar and remembered objects, with more evidence for recollection from ca. 500 ms onward.

It is an open question whether the neural signals for memory representations and decision-making processes play a causal role in supporting mnemonic experiences, such as familiarity and recollection. Additionally, it is not well understood how these signals manifest in the case of false memories. To date, reliable neural markers of false memories have not been established (Xue, 2018).

This present study made use of the flexibility of MVPA to explore these questions by selecting and relabeling evoked responses in the training datasets (see **Supplementary Information Table 1B**) to align with those of the test experiment. In the case of object-scene associations (where no novel items were presented in the test phase), this was achieved by changing the “remembered” and “forgotten” labels to “familiar” and “unfamiliar”. In the case of objects, for false alarms, this meant relabeling novel (i.e., first presented in the test phase) objects for which participants responded “familiar” or “remembered” as such, while keeping the “unfamiliar” labels of correctly identified novel objects. Remarkably, training remembered and forgotten object/scene pairs, and on false alarms for both familiar and remembered objects, yielded robust and sustained cross-classification patterns.

For false-alarm-familiar objects the patters of cross-classification accuracies were very similar to those of correctly identified familiar objects up to ca. 600 ms, with an extended positive decoding effect (**Figure 6a**). In contrast to correctly identified remembered objects, training on false-alarm-remembered objects (**Figure 6b**) led to early effects as well; the steeply rising late-phase peak, while present, was delayed by ca. 20-50 ms over central-posterior ROIs (see **Supplementary Information 5**). In the case of forgotten (versus novel) objects, cross-classification over all electrodes yielded a positive cluster for the 400 to 600 ms window, while the early peak seen in the case of familiar objects in bilateral posterior and right central regions of interest was not observed, and no significant clusters were found in frontal ROIs.

### Are faces ‘special’?

The human face is possibly the most extensively investigated high-level visual stimulus in cognitive science, and face perception is often viewed as a model system for investigating the development and organization of information processing in the brain (Scherf et al., 2012; Young, 2018; Yovel et al., 2014). The social and personal importance, as individuals and as a species, makes faces a salient stimulus category. Due to some of the very same reasons, the generalizability of the finding in face perception research to other stimulus categories is a matter of debate, summarized in the question: “are faces special?” (Liu and Chaudhuri, 2003; Yue et al., 2006). In the field of memory research, arguments are made that the recognition of faces may be “inherently different from recognition of other kinds of nonverbal stimuli” (Danker et al., 2008). The question whether the processes that contribute to face recognition are unique can be probed by investigating how these processes unfold for various other stimulus types (Tovée, 1998). In this present study, the choice of a face recognition dataset as the test set makes it possible to tackle this question.

The results presented here show substantial overlap between recognition signals for faces and other stimulus categories. These effects were most consistent in the later, ca. post-300-400 ms phase, but were also found in the earlier, 200 ms time window in some cases (see the previous **Early and Late Effects** section). Training classifiers on ERPs for various stimulus types and using these to test familiarity in ERPs for highly personally familiar and unfamiliar faces yielded extensive positive effects, arguing for similar information processing.

On first sight, the results of this analysis argue for at least a partial overlap between systems that subserve the recognition of faces and other stimulus categories, particularly in the later (ca. 400 ms onward) stages of processing. However, alternative explanations for this shared effect must be considered. Due to the low spatial resolution of EEG, the exact structural localization of the origin of these signals is challenging. For example, regions thought to be category-selective are found in nearby anatomical locations in the lateral occipital (e.g., OFA for faces, LO for objects, EBA for bodies, OPA for places) and temporal cortices (FFA for faces, pFs for objects, FBA for bodies, PPA for scenes) (Kamps et al., 2016; Kanwisher, 2010; Taylor et al., 2007; Weiner et al., 2018). It is thus possible that while successful cross-classification in part reflects similar activity in the same areas of the brain (e.g., the hippocampus), other parts of the shared signal may result from information flow in parallel processing pathways between different nodes within centimeters of distance. Source-localized MEG and fMRI-M/EEG fusion methods should investigate this question in the future.

### Time since acquisition

Three of the experiments in this report used single-session experimental familiarization: Sommer et al. (2021) involved a 10 to 60 s familiarization phase with a previously unfamiliar single face image immediately before the start of the experiment. The 180-item acquisition phase and the test phase in Dimsdale-Zucker et al. (2022) was separated by a 45–60-minute interval. Treder et al. (2021) had eight, 32-item encoding/recall blocks, each with a 3-minute delay between learning and test. In two studies, stimuli were pre-experimentally familiar: Wakeman and Henson (2015) used faces of well-known celebrities, while Jagiello et al. (2019) presented personally relevant music. Stimuli in the test dataset (Wiese et al., 2022) were also pre-experimentally familiar (faces of personally highly familiar identities). Despite these differences, consistent and overlapping cross-classification accuracies were observed in all analyses.

Old/new effects similar to those observed for long-term memory have been observed for short-term memory tasks. Using a wide range of stimulus types (single letters, words and pseudowords, pictures of objects, dots at different spatial locations, and two-dimensional sinusoidal gratings), Danker and colleagues (Danker et al., 2008) have found that the amplitude of the FN400 showed old/new effects only for verbal stimuli, which increased with recency. Old/new effects indexed by the LPC, on the other hand, were seen for a range of stimulus types, and were not modulated by recency.

Although systematic studies on other stimulus types are yet to be conducted, EEG patterns for short-term, experimentally familiarized faces have been shown to generalize to longer-term personal familiarity (Dalski et al., 2022a, 2022b). The results of this present study further support these findings, as training on single-session familiarization data, in addition to faces (Sommer et al., 2021), familiar objects (Dimsdale-Zucker et al., 2022) and object-scene associations (Treder et al., 2021), all yielded positive effects.

### Negative decoding

Post-500 ms sustained, significant below-chance classification accuracies were seen in several training datasets. Similar below-chance classification patterns are commonly observed, especially in temporal generalization analyses (Carlson et al., 2013, 2011; King et al., 2014; Nikolić et al., 2009). Interestingly, the reasons for such reversal phenomena are not yet understood (King and Dehaene, 2014), and the issue has received little attention so far. That this reversal is not merely an epiphenomenon related to the end of the stimulus presentation but is a meaningful effect that reflects cognitive processing is indicated by its attenuation when false alarms are tested against correct rejections in the familiar/remembered objects experiment (**Figure 6**A and B, **Supplementary Information 5**). Sustained positive cross-decodability in these cases may reflect neural activity related to extended search processes, or, alternatively, extra-experimental familiarity.

In this present investigation, sustained, significant below-chance classification performance was only observed in studies that involved short-term, laboratory-based familiarization methods, i.e., in the cases of the experimentally familiarized face and the familiar/remembered objects experiments. Cross-classification did not yield such patterns when the pre-experimentally familiar (famous) faces and pre-experimentally familiar music datasets were used for training. Thus, it is possible that such pattern reversal effects are related to control processes for items with recent or weak memory traces.

Further complicating the interpretation of these findings is the fact that the studies where stimuli were newly learned were also the ones that directly tested recollection for these items. The experimentally familiarized face and the familiar/remembered objects experiment both required responses that pertained to memory recall. The familiar music experiment required no response, while the task – symmetry judgement – was not related to familiarity in the famous faces study. As the type of the required response and the length of familiarity were not independent factors, further research is needed to study the causes of this phenomenon.

### Further considerations

This analysis was based on publicly available EEG datasets that were collected in different laboratories using different EEG systems under varying conditions. The stimulus presentation times, number of trials and repetitions, and control for low-level properties also varied significantly across the studies. In addition to the stimulus categories analyzed in this study, some of the experiments also presented additional stimulus types, such as the participant’s own face in the study by Sommer et al. (2021), scrambled faces in the study by Wakeman & Henson (2015), and a “no variance” condition using a second familiar/unfamiliar pair in the study by Wiese et al. (2022). It is highly likely that the complete stimulus pool has an influence on the neural signals for individual stimulus conditions. Therefore, it is important to note that this analysis does not represent a systematic investigation into the factors that contribute to successful cross-classification. Further research is needed to systematically investigate these factors, and to replicate and extend these findings in more controlled and standardized conditions.

Considering that the spatio-temporal pattern of the effects is dependent on the information content in both the training and test datasets, neural signals of recognition that are not present in either the training or test dataset will not be captured by the analysis. Therefore, it is important to keep in mind that the current study used a single test dataset that focused on face familiarity. Future studies should explore the potential effects of other stimulus types, such as written and spoken words, which are commonly used in neuroscience research and have not been investigated here. Additionally, it would be interesting to explore how other sensory modalities, such as auditory or tactile stimuli, may influence these neural representations. Furthermore, it would be valuable to examine the interplay between recollection, neural representations of various stimulus properties, and attention in the context of familiarity and recollection.

## Conclusion

In this study, multivariate cross-dataset classification analysis was used to investigate the shared signals of recognition memory across a range of different stimulus types and experimental conditions. This approach allows for the identification of patterns of neural activity that are shared across different conditions and datasets, providing a more comprehensive view of the neural dynamics underlying memory processes. The results indicate that the neural signals for face stimuli generalized remarkably well starting from around 200 ms following stimulus onset. For non-face stimuli, successful cross-classification was most robust in the later period. Furthermore, a dissociation between familiar and remembered objects was observed, with shared signals present only in the late window for correctly remembered objects, while cross-decoding results for familiar objects were present in the early period as well. Furthermore, the subjective sense of familiarity and/or recollection, i.e., false alarms and incorrect responses, also produced similar patterns of cross-classification. Overall, these findings show that multivariate cross-classification across datasets can provide valuable insights into the neural dynamics of memory processes.

## Data and code availability

Data and code will be made openly available.

## Ethics statement

The study was approved by the Bournemouth University Research Ethics committee (ID 46391).

### Acknowledgments

This research received no specific grant from any funding agency in the public, commercial, or not-for-profit sectors.

## Author contributions

GGA is the sole author of the manuscript.

## Supplementary Materials

**Supplementary Information**. Extended methods and results

**Supplementary Table 1**. Statistics for the main time-resolved classification analyses. All sensors and pre-defined regions of interest

**Supplementary Table 2**. Statistics for the additional analyses conducted on the *objects* dataset

**Supplementary Table 3**. Statistics for the main temporal generalization analyses. All sensors and pre-defined regions of interest

**Supplementary Table 4**. Statistics for the additional temporal generalization analyses conducted on the *objects* dataset

**Supplementary Table 5**. Statistics for the main spatiotemporal searchlight analyses

**Supplementary Table 6**. Statistics for the additional spatiotemporal searchlight analyses conducted on the *objects* dataset

