## Supplementary Information for "Shared neural codes of recognition memory"

### Supplementary Information 1

Cross-classification was performed on common channels in the training datasets and test dataset. These common channels are presented in Figure S1.

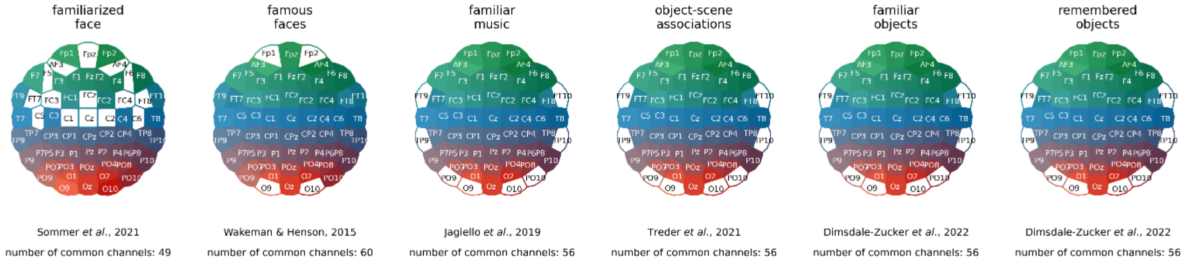


**Supplementary Information, Figure S1.** Shared EEG channels between the test dataset (Wiese et al., 2022) and the training datasets (Dimsdale-Zucker et al., 2022; Jagiello et al., 2019; Sommer et al., 2021; Treder et al., 2021; Wakeman and Henson, 2015). Supplements the *Methods* section of the main text.

Tale S1A gives an overview of the datasets included in the study, such as number of participants, experimental tasks, stimulus presentation times, and number of trials. In some experiments, additional stimulus categories were also presented, which were not analyzed in this present study.

| | Dataset | Study | *n* | Task | Presentation time/ *n* trials | Stimuli/conditions not included |
|---|---|---|---|---|---|---|
| TEST | personally familiar and unfamiliar faces | Wiese *et al.* (2021) | 22 | *No task* for study-relevant items. *Not stimulus-related*: butterfly-detection task for maintaining attention | 1000 ms<br>50/50 trials | One additional familiar and unfamiliar identity with a single image |
| TRAINING | experimentally familiarized face and unfamiliar faces | Sommer *et al.* (2021) | 15 | *Stimulus-related*: Detecting the familiarized face | 500 ms<br>792 trials | Own face |
| | famous and unfamiliar faces | Wakeman & Henson (2015) | 15 | *Stimulus-related*: Symmetry judgement | 800 to 1000 ms<br>600 trials † | Scrambled faces |
| | familiar and unfamiliar music | Jagiello *et al.* (2019) | 10 | *No task* – passive listening to short segments of pre-experimentally familiar and unfamiliar music | 750 ms<br>200 trials ‡ | - |
| | remembered and forgotten object-scene associations | Treder *et al.* (2021) | 20 | *Stimulus-related*: indicate if the paired associate of stimulus is remembered or forgotten | 2500 to 6000 ms<br>256 trials | |
| | familiar and novel objects | Dimsdale-Zucker *et al.* (2022) | 38 | *Stimulus-related*: “remembered”, “familiar”, or “new” decision | 700 ms | - |
| | remembered and novel objects | | | | 270 trials | - |

**Supplementary Information, Table S1A.** An overview on the datasets analyzed in the study. Supplements the *Methods* section in the main text. The number of trials is the nominal number of presentations for all stimulus types/conditions included in the analyses.
*Notes*: † In the *famous and unfamiliar faces* experiment, trials in which famous faces were not recognized, and unfamiliar faces that were falsely indicated as familiar, were not analyzed. ‡ Only averaged ERPs (i.e., 1 familiar and 1 unfamiliar sample per participant) were available in the *familiar and unfamiliar music* experiment.

Training labels can be changed to test cross-classification performance across different tasks and domains (Dalski et al., 2022). This study made use of this flexibility of the MVPA method to probe different aspects of memory functions. Table 1B gives an overview on how labels were modified for this purpose.

| | Dataset | Study | Original condition | New label | Data included |
|---|---|---|---|---|---|
| TEST | personally familiar and unfamiliar faces | Wiese *et al.* (2021) | Face-images of a personally familiar person | Familiar | Data: 22 participants 44.95±4.7 trials: min: 30, max: 50 |
| | | | Face-images of an unfamiliar person | Unfamiliar | |
| TRAINING | experimentally familiarized face and unfamiliar faces | Sommer *et al.* (2021) | An experimentally familiarized image of a person | Familiar | Data: 15 participants 65.66±9.7 trials: min: 33, max: 72 |
| | | | Images of unfamiliar persons | Unfamiliar | |
| | famous and unfamiliar faces | Wakeman & Henson (2015) | Face-images of famous individuals | Familiar | Data: 15 participants 226.6±45.7 trials: min: 148, max: 292 |
| | | | Face-images of unfamiliar individuals | Unfamiliar | |
| | familiar and unfamiliar music | Jagiello *et al.* (2019) | Personally familiar and relevant music | Familiar | Averaged ERPs, one per condition per participant |
| | | | Unfamiliar music | Unfamiliar | |
| | remembered and forgotten object-scene associations | Treder *et al.* (2021) | Remembered object/scene associations | Familiar | Data: 18 participants 46.5±16.8 trials: min: 21, max: 77 |
| | | | Forgotten object/scene associations | Unfamiliar | |
| | familiar and novel objects | Dimsdale-Zucker *et al.* (2022) | Old items correctly categorized as familiar | Familiar | Data: 38 participants 55.97±14.5 trials: min: 32, max: 86 |
| | | | New items correctly categorized as new | Unfamiliar | |
| | remembered and novel objects | | Old items correctly categorized as remembered | Familiar | Data: 38 participants 62.84±16.0 trials: min: 31, max: 106 |
| | | | New items correctly categorized as new | Unfamiliar | |
| | False alarms for familiar objects | | New items incorrectly categorized as familiar | Familiar | Data: 38 participants 15.92±9.9 trials: min: 2, max: 48 |
| | | | New items correctly categorized as new | Unfamiliar | |
| | False alarms for remembered objects | | New items incorrectly categorized as remembered | Familiar | Data: 24 participants 4.70±5.0 trials: min: 1, max: 19 |
| | | | New items correctly categorized as new | Unfamiliar | |
| | Forgotten objects | | Old items incorrectly categorized as new | Familiar | Data: 38 participants 35.02±17.7 trials: min: 6, max: 77 |
| | | | New items correctly categorized as new | Unfamiliar | |
| | Subjective recollection for incorrect responses | | Old items incorrectly categorized as new | Familiar | Data: 38 participants 14.74±9.9 trials: min: 3, max: 58 |
| | | | New items incorrectly categorized as old | Unfamiliar | |
| | Familiarity vs. recollection | | Old items correctly categorized as familiar | Familiar | Data: 38 participants 51.39±12.75 trials: min: 31, max: 82 |
| | | | Old items correctly categorized as remembered | Remembered | |

**Supplementary Information, Table S1B. Relabeling the conditions for cross-dataset train-test procedures.**
Training and testing were performed on balanced data, under-sampling to the minimum trial count available for the given participant. The 'data included' column presents the number of participants included in the analysis, the mean ± SD number of trials per participant, and the minimum and maximum number of trials for individual participants. Supplements the *Methods* section in the main text.

### Supplementary Information 2

For visualization purposes, the results shown in **Figure 3** in the main text are presented in here, in **Figure S2**, in an alternative format. For ease of comparison, time-resolved cross-classification involving face datasets are presented on the right, the music and object/scene associations datasets in the middle, and the correctly remembered and familiar objects on the right.

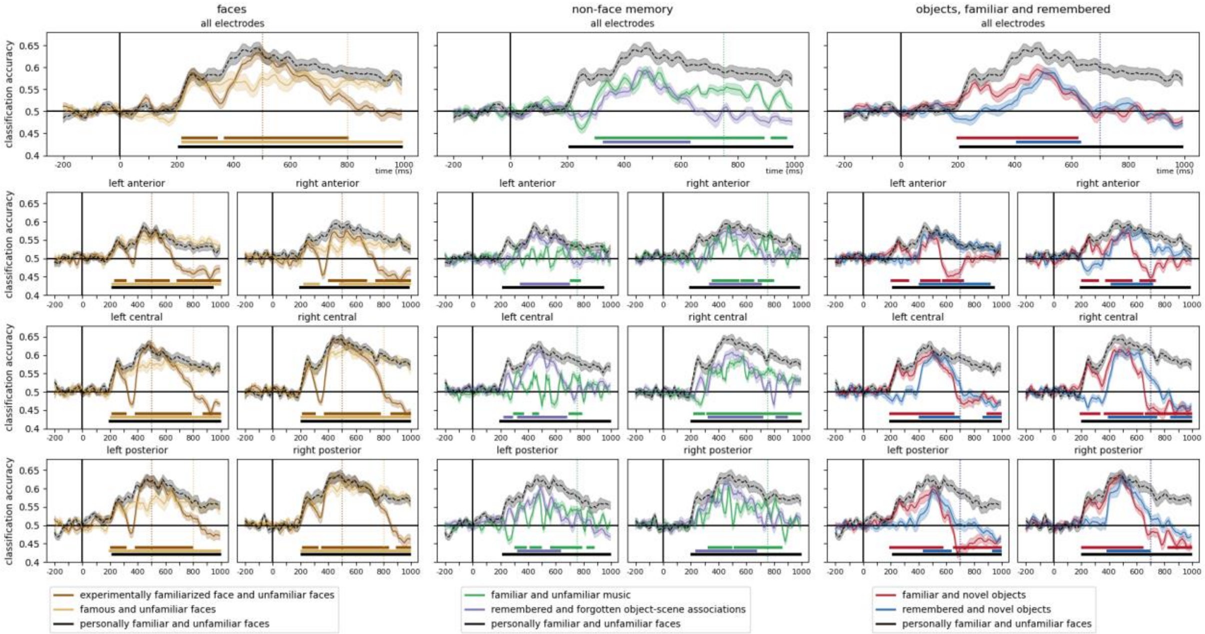


**Supplementary Information, Figure S2.** Time-resolved cross-classification, supplements **Figure 3** in the main text. Cross classification accuracies for face datasets are presented on the right, the remaining music and object/scene associations in the middle, and familiar and remembered objects on the right. Leave-one-subject-out classification accuracies for the test dataset are overlayed in black. Shaded regions denote ±SEM. Horizontal lines denote significant clusters (two-sided cluster permutation tests, $p<0.05$). **Supplementary Table 1**. presents detailed statistics.

### Supplementary Information 3

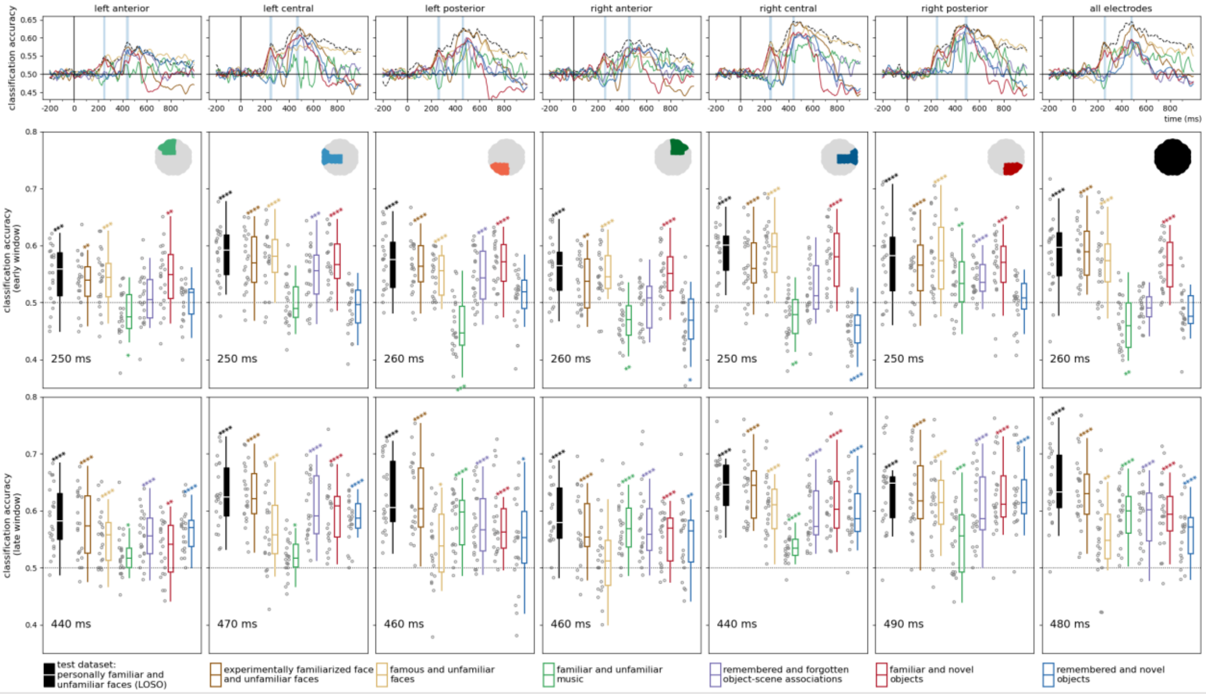


**Supplementary Information, Figure S3A. Classification accuracies in the early and late phases in the pre-defined regions of interest, and over all electrodes.** Ealy and late time-points correspond to peak leave-one-subject-out classification accuracies between 200 and 270 ms (early) and 400 and 1000 ms (late) in the test dataset (indicated in the lower-left corner in each panel).

Statistics: two-sided one-sample *t*-tests, $p_{uncorrected}$ < 0.05*, <0.01**, <0.001***, <0.0001****; boxplots: first, second (median), and third quartiles; whiskers with Q1 − 1.5 IQR and Q3 + 1.5 IQR, with horizontally jittered individual classification accuracies. Supplements **Figure 4** in the main text.

Most prominent effects were observed in the right posterior and central clusters, and **Figure 4** in the main text shows early and late effects in detail for these regions of interest. Cross-classification accuracy in early and late phases, in all regions of interest are presented in **Figure S3A.** Cross-classification accuracies in all regions of interest, averaged in 100 ms bins, are presented in **Figure S3B**.

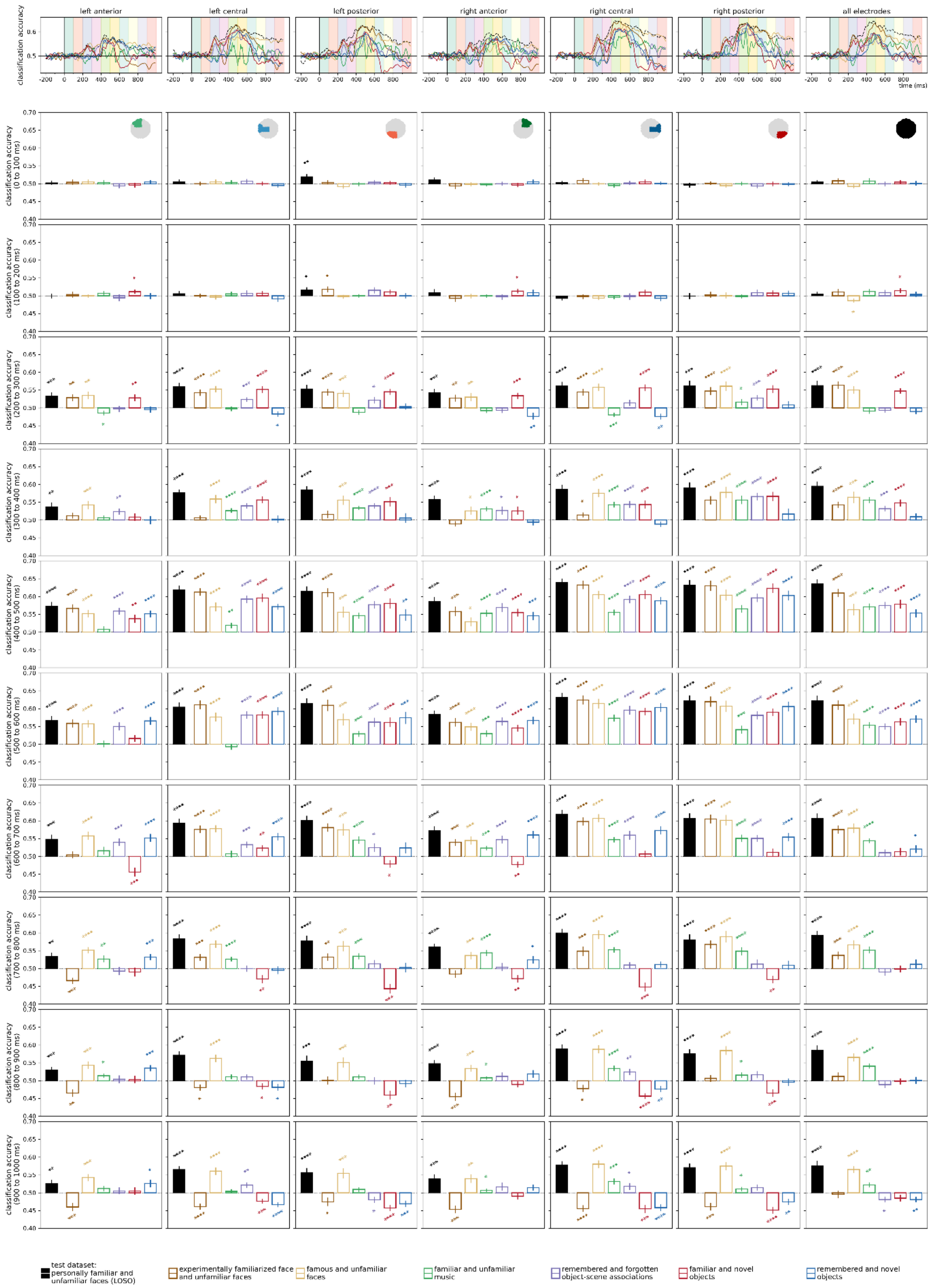


**Supplementary Information, Figure S3B. Classification accuracy scores averaged over 100 ms bins in the pre-defined regions of interest, and over all electrodes.** Bar charts and error ranges represent mean ±SEM. Statistics: two-sided one-sample *t*-tests against chance, $p_{uncorrected}$ < 0.05*, <0.01**, <0.001***, <0.0001****

**Supplementary Information 4**

In their original report, Jagiello and colleagues (2019) noted observing an early familiarity effect in pupillary responses starting around ca. 100 ms, for which a corresponding early ERP component was not seen. Here, in the interest of completeness, cross-classification accuracy scores, averaged between the training times 100 and 170 ms, are presented. Significant, sustained positive clusters are found in central and posterior clusters, starting around 270-330 ms, while negative or no effects are seen in anterior clusters and over all electrodes.

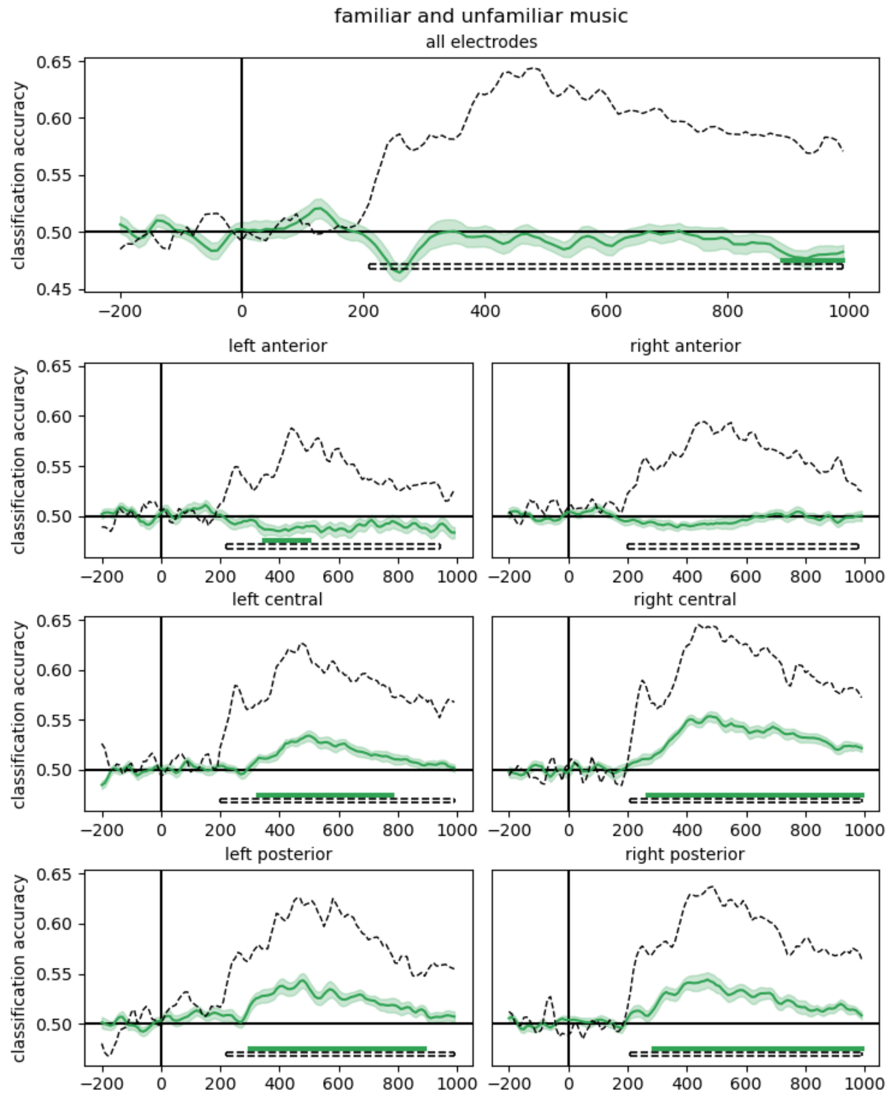


all electrodes 890 to 990 ms, cluster p = 0.0264

| | | | |
|---|---|---|---|
| left anterior | 350 to 500 ms, cluster p = 0.0189 | right anterior | - |
| left central | 330 to 780 ms, cluster p = 0.0004 | right central | 270 to 990 ms, cluster p = 0.0001 |
| left posterior | 300 to 890 ms, cluster p = 0.0001 | right posterior | 290 to 990 ms, cluster p = 0.0001 |

**Supplementary Information, Figure S4**. Trained on familiar and unfamiliar music, tested on personally familiar and unfamiliar faces. Classification accuracies between 100 and 170 ms training times are averaged. The results of the leave-one-subject-out classification for familiar faces is presented as dashed line for comparison. Shaded regions denote ±SEM. Horizontal lines denote significant clusters (two-sided cluster permutation tests, $p<0.05$).

### Supplementary Information 5.

Supplementary Figure S5 presents the results of **Figure 6A** and **B** in the main text, depicting the results of cross-classification on false alarms, in an alternative format. For ease of comparison, results for correctly identified familiar/remembered objects (also shown in **Figure 3**, and **Figure S2**) are presented here as well.

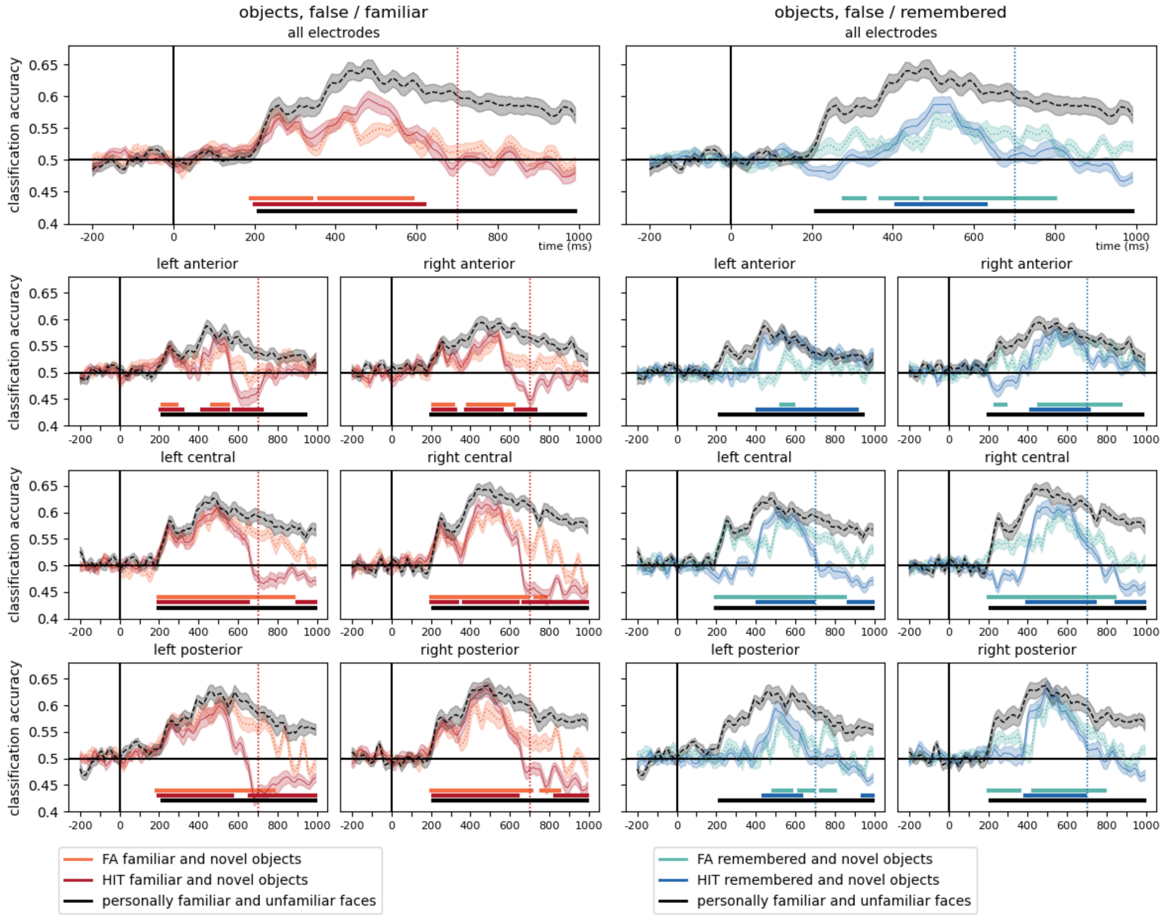


**Supplementary Information, Figure SX.** The time-course of cross-classification accuracies for familiar/remembered objects, when training was performed on false alarms (novel objects, incorrectly categorized as either familiar or remembered) and correct rejections (novel objects correctly identified as new). The results of the within-experiment LOSO for the test dataset, and the hits and correct rejections cross-classification, are also presented for comparison. Two-sided cluster permutation tests ($p$<0.05), shaded ranges denote ±SEM. For detailed statistics, see **Supplementary Table 2**.
