## Supplementary Table for "Shared neural codes of recognition memory": Supplementary_Table_1.html

  

### Supplementary Table 1.

  

Results of the main time-resolved classification analyses

  

A) **personally familiar and unfamiliar faces** (leave-one-subject-out)

|  |  |  |  |  |  |  |  |  |
| --- | --- | --- | --- | --- | --- | --- | --- | --- |
|  | time window | peak latency | cluster *p* | peak Cohen's *d* |  | | | |
| **all electrodes** | 210 - 990 ms | 480 ms | 0.0001 | 2.1658 |  | | | |
|  | **left hemisphere** | | | | **right hemisphere** | | | |
|  | time window | peak latency | cluster *p* | peak Cohen's *d* | time window | peak latency | cluster *p* | peak Cohen's *d* |
| **anterior** | 220 - 940 ms | 440 ms | 0.0001 | 1.575 | 200 - 980 ms | 460 ms | 0.0001 | 1.6355 |
| **central** | 200 - 990 ms | 470 ms | 0.0001 | 2.242 | 210 - 990 ms | 440 ms | 0.0001 | 2.9577 |
| **posterior** | 220 - 990 ms | 460 ms | 0.0001 | 2.016 | 210 - 990 ms | 490 ms | 0.0001 | 1.9183 |

  

B) **experimentally familiarized face and unfamiliar faces**

|  | time window | peak latency | cluster *p* | peak Cohen's *d* |  | | | |  |  |
| **all electrodes** | 220 - 340 ms | 250 ms | 0.0131 | 1.6881 |  | | | |  |  |
 370 - 800 ms | 490 ms | 0.0001 | 1.9626 |  | | | ||  |  | | | | | | | | |  |
|  | **left hemisphere** | | | | **right hemisphere** | | | |  |  |
|  | time window | peak latency | cluster *p* | peak Cohen's *d* | time window | peak latency | cluster *p* | peak Cohen's *d* |  |  |
| **anterior** | 240 - 310 ms | 260 ms | 0.037 | 0.9389 | 410 - 670 ms | 540 ms | 0.0001 | 1.0937 |  |  |
 390 - 620 ms | 450 ms | 0.0001 | 1.2688 | 750 - 990 ms | 750 ms | 0.0008 | -0.5387 | 690 - 990 ms | 690 ms | 0.0004 | -0.4455 |  | | | ||  | **central** | 220 - 310 ms | 250 ms | 0.0258 | 1.3608 | 220 - 300 ms | 260 ms | 0.0281 | 1.4837 |  |
 390 - 780 ms | 480 ms | 0.0001 | 2.2827 | 380 - 770 ms | 490 ms | 0.0001 | 2.2552 | 880 - 990 ms | 880 ms | 0.0213 | -0.5059 | 860 - 990 ms | 860 ms | 0.0198 | -0.6595 ||  |  | **posterior** | 210 - 310 ms | 260 ms | 0.0225 | 1.3659 | 220 - 320 ms | 260 ms | 0.0283 | 1.1184 |
 390 - 790 ms | 470 ms | 0.0001 | 1.7943 | 360 - 830 ms | 470 ms | 0.0001 | 2.0643 |  | | | | 900 - 990 ms | 900 ms | 0.043 | -0.5385 |

  

C) **famous and unfamiliar faces**

|  | time window | peak latency | cluster *p* | peak Cohen's *d* |  | | | |  |
| **all electrodes** | 220 - 990 ms | 640 ms | 0.0002 | 1.4114 |  | | | |  |
|  | | | | | | | | |  |
|  | **left hemisphere** | | | | **right hemisphere** | | | |  |
|  | time window | peak latency | cluster *p* | peak Cohen's *d* | time window | peak latency | cluster *p* | peak Cohen's *d* |  |
| **anterior** | 220 - 990 ms | 620 ms | 0.0001 | 1.3066 | 230 - 330 ms | 260 ms | 0.0269 | 1.0276 |  |
  | | | | 490 - 990 ms | 540 ms | 0.0001 | 1.1556 ||  | **central** | 210 - 990 ms | 600 ms | 0.0001 | 1.9211 | 210 - 990 ms | 540 ms | 0.0001 | 1.8772 |
| **posterior** | 200 - 990 ms | 640 ms | 0.0002 | 1.2837 | 210 - 990 ms | 540 ms | 0.0001 | 1.4273 |  |

  

D) **familiar and unfamiliar music**

|  | time window | peak latency | cluster *p* | peak Cohen's *d* |  | | | |  |  |
| **all electrodes** | 300 - 890 ms | 470 ms | 0.0001 | 2.0639 |  | | | |  |  |
 920 - 970 ms | 950 ms | 0.0351 | 1.2991 |  | | | ||  |  | | | | | | | | |  |
|  | **left hemisphere** | | | | **right hemisphere** | | | |  |  |
|  | time window | peak latency | cluster *p* | peak Cohen's *d* | time window | peak latency | cluster *p* | peak Cohen's *d* |  |  |
| **anterior** | 710 - 770 ms | 740 ms | 0.0114 | 1.0492 | 360 - 540 ms | 470 ms | 0.0002 | 1.5712 |  |  |
  | | | | 570 - 650 ms | 610 ms | 0.0103 | 0.8743 |  | | | | 690 - 730 ms | 720 ms | 0.0134 | 1.5837 |  | | | | 750 - 790 ms | 770 ms | 0.0179 | 1.5338 ||  | **central** | 300 - 360 ms | 330 ms | 0.0005 | 1.1508 | 320 - 990 ms | 580 ms | 0.0001 | 1.8132 |  |
 440 - 470 ms | 440 ms | 0.0087 | 1.3923 | 230 - 290 ms | 290 ms | 0.0389 | -0.467 | 700 - 780 ms | 730 ms | 0.0002 | 1.2877 |  | | | ||  |  | **posterior** | 310 - 380 ms | 340 ms | 0.0043 | 1.6852 | 330 - 490 ms | 470 ms | 0.0015 | 1.5293 |
 420 - 490 ms | 470 ms | 0.0036 | 1.6951 | 520 - 770 ms | 590 ms | 0.0009 | 1.0399 | 570 - 780 ms | 590 ms | 0.0003 | 1.0363 | 790 - 850 ms | 810 ms | 0.015 | 1.4681 | 830 - 870 ms | 830 ms | 0.0363 | 0.9979 |  | | | |

  

E) **remembered and forgotten object-scene associations**

|  | time window | peak latency | cluster *p* | peak Cohen's *d* |  | | | |  |
| **all electrodes** | 330 - 630 ms | 450 ms | 0.0001 | 2.2479 |  | | | |  |
|  | | | | | | | | |  |
|  | **left hemisphere** | | | | **right hemisphere** | | | |  |
|  | time window | peak latency | cluster *p* | peak Cohen's *d* | time window | peak latency | cluster *p* | peak Cohen's *d* |  |
| **anterior** | 350 - 690 ms | 460 ms | 0.0001 | 1.6205 | 340 - 700 ms | 460 ms | 0.0001 | 1.1367 |  |
| **central** | 230 - 280 ms | 260 ms | 0.0367 | 1.057 | 330 - 710 ms | 500 ms | 0.0001 | 1.7539 |  |
 330 - 670 ms | 490 ms | 0.0001 | 1.7813 | 820 - 890 ms | 840 ms | 0.0428 | 0.8474 ||  | **posterior** | 330 - 630 ms | 490 ms | 0.0001 | 1.5454 | 240 - 670 ms | 470 ms | 0.0001 | 1.7491 |

  

F) **familiar and novel objects**

|  | time window | peak latency | cluster *p* | peak Cohen's *d* |  | | | |  |  |
| **all electrodes** | 200 - 620 ms | 480 ms | 0.0001 | 1.5964 |  | | | |  |  |
|  | | | | | | | | |  |  |
|  | **left hemisphere** | | | | **right hemisphere** | | | |  |  |
|  | time window | peak latency | cluster *p* | peak Cohen's *d* | time window | peak latency | cluster *p* | peak Cohen's *d* |  |  |
| **anterior** | 210 - 320 ms | 250 ms | 0.0203 | 0.8119 | 210 - 320 ms | 250 ms | 0.0113 | 1.2088 |  |  |
 420 - 550 ms | 480 ms | 0.0031 | 1.1423 | 380 - 560 ms | 500 ms | 0.0001 | 1.2355 | 580 - 720 ms | 720 ms | 0.0051 | -0.5117 | 630 - 730 ms | 630 ms | 0.012 | -0.4439 ||  | **central** | 200 - 650 ms | 490 ms | 0.0001 | 1.8137 | 200 - 330 ms | 250 ms | 0.0095 | 1.3771 |  |
 900 - 990 ms | 930 ms | 0.0404 | -0.4573 | 370 - 640 ms | 480 ms | 0.0001 | 1.8587 |  | | | | 670 - 990 ms | 840 ms | 0.0002 | -0.7578 ||  |  | **posterior** | 200 - 570 ms | 500 ms | 0.0001 | 1.5908 | 210 - 640 ms | 480 ms | 0.0001 | 2.0162 |
 660 - 990 ms | 660 ms | 0.0002 | -0.6514 | 830 - 990 ms | 830 ms | 0.009 | -0.4528 |

  

G) **remembered and novel objects**

|  | time window | peak latency | cluster *p* | peak Cohen's *d* |  | | | |  |
| **all electrodes** | 410 - 630 ms | 540 ms | 0.0001 | 1.55 |  | | | |  |
|  | | | | | | | | |  |
|  | **left hemisphere** | | | | **right hemisphere** | | | |  |
|  | time window | peak latency | cluster *p* | peak Cohen's *d* | time window | peak latency | cluster *p* | peak Cohen's *d* |  |
| **anterior** | 410 - 910 ms | 540 ms | 0.0001 | 1.3653 | 420 - 710 ms | 540 ms | 0.0001 | 1.5775 |  |
| **central** | 410 - 690 ms | 520 ms | 0.0001 | 1.7662 | 400 - 740 ms | 490 ms | 0.0001 | 1.7627 |  |
 870 - 990 ms | 870 ms | 0.0169 | -0.5517 | 850 - 990 ms | 850 ms | 0.0111 | -0.5043 ||  | **posterior** | 440 - 630 ms | 500 ms | 0.0055 | 1.3535 | 390 - 690 ms | 500 ms | 0.0001 | 1.9123 |
 940 - 990 ms | 940 ms | 0.048 | -0.6084 |  | | | |

  
