## Supplementary Table for "Shared neural codes of recognition memory": Supplementary_Table_2.html

  


### Supplementary Table 2.

  

Results of the time-resolved classification analyses, trained on the familiar - remembered object dataset

  

A) **FA familiar and novel objects**

|  | time window | peak latency | cluster *p* | peak Cohen's *d* |  | | | |  |
| **all electrodes** | 190 - 340 ms | 260 ms | 0.0006 | 1.4865 |  | | | |  |
 360 - 590 ms | 430 ms | 0.0001 | 1.6413 |  | | | ||  |  | | | | | | | | |
|  | **left hemisphere** | | | | **right hemisphere** | | | |  |
|  | time window | peak latency | cluster *p* | peak Cohen's *d* | time window | peak latency | cluster *p* | peak Cohen's *d* |  |
| **anterior** | 220 - 290 ms | 250 ms | 0.0401 | 0.9053 | 210 - 310 ms | 250 ms | 0.0284 | 0.8732 |  |
 470 - 550 ms | 530 ms | 0.0253 | 0.8579 | 390 - 620 ms | 540 ms | 0.0029 | 1.4097 ||  | **central** | 200 - 880 ms | 480 ms | 0.0001 | 2.2229 | 200 - 690 ms | 490 ms | 0.0001 | 1.7556 |
  | | | | 730 - 780 ms | 750 ms | 0.0465 | 1.2953 || **posterior** |  | 190 - 780 ms | 560 ms | 0.0001 | 1.5545 | 200 - 710 ms | 420 ms | 0.0003 | 1.4831 |
  | | | | 760 - 850 ms | 830 ms | 0.0349 | 1.2088 |

  

B) **FA remembered and novel objects**

|  | time window | peak latency | cluster *p* | peak Cohen's *d* |  | | | |  |
| **all electrodes** | 280 - 330 ms | 310 ms | 0.0308 | 1.3011 |  | | | |  |
 370 - 460 ms | 420 ms | 0.0155 | 0.9395 |  | | | | 480 - 800 ms | 510 ms | 0.0001 | 1.2878 |  | | | ||  |  | | | | | | | | |
|  |  | **left hemisphere** | | | | **right hemisphere** | | | |
|  | time window | peak latency | cluster *p* | peak Cohen's *d* | time window | peak latency | cluster *p* | peak Cohen's *d* |  |
| **anterior** | 530 - 590 ms | 550 ms | 0.0471 | 0.6888 | 240 - 290 ms | 260 ms | 0.0453 | 1.1206 |  |
  | | | | 460 - 780 ms | 550 ms | 0.0001 | 1.5254 |  | | | | 800 - 870 ms | 820 ms | 0.0308 | 0.8444 ||  | **central** | 200 - 850 ms | 520 ms | 0.0001 | 2.0578 | 200 - 840 ms | 560 ms | 0.0001 | 1.493 |
|  | **posterior** | 490 - 580 ms | 520 ms | 0.005 | 1.1208 | 200 - 360 ms | 350 ms | 0.0099 | 0.7633 |
 620 - 690 ms | 630 ms | 0.0168 | 1.0377 | 430 - 790 ms | 540 ms | 0.0001 | 1.478 | 730 - 800 ms | 750 ms | 0.0085 | 1.0906 |  | | | |

  

C) **forgotten vs. novel objects**

|  | time window | peak latency | cluster *p* | peak Cohen's *d* |  | | | |  |
| **all electrodes** | 370 - 560 ms | 490 ms | 0.0056 | 1.1985 |  | | | |  |
|  | | | | | | | | |  |
|  | **left hemisphere** | | | | **right hemisphere** | | | |  |
|  | time window | peak latency | cluster *p* | peak Cohen's *d* | time window | peak latency | cluster *p* | peak Cohen's *d* |  |
| **anterior** |  | | | |  | | | |  |
| **central** | 200 - 780 ms | 470 ms | 0.0001 | 2.2515 | 290 - 770 ms | 490 ms | 0.0001 | 1.9893 |  |
 800 - 850 ms | 820 ms | 0.0495 | 1.0298 |  | | | ||  | **posterior** | 280 - 680 ms | 450 ms | 0.0001 | 1.6312 | 260 - 760 ms | 510 ms | 0.0001 | 1.9095 |

  

D) **objects, incorrect answers with subjective labels**

|  | time window | peak latency | cluster *p* | peak Cohen's *d* |  | | | |  |
| **all electrodes** | 210 - 270 ms | 250 ms | 0.0262 | 1.3828 |  | | | |  |
 360 - 450 ms | 390 ms | 0.0059 | 1.4467 |  | | | | 470 - 610 ms | 580 ms | 0.0041 | 1.1294 |  | | | ||  |  | | | | | | | | |
|  |  | **left hemisphere** | | | | **right hemisphere** | | | |
|  | time window | peak latency | cluster *p* | peak Cohen's *d* | time window | peak latency | cluster *p* | peak Cohen's *d* |  |
| **anterior** | 380 - 440 ms | 430 ms | 0.035 | 1.131 | 380 - 610 ms | 520 ms | 0.0001 | 1.1033 |  |
| **central** | 360 - 450 ms | 420 ms | 0.0154 | 1.3904 | 360 - 880 ms | 580 ms | 0.0001 | 2.1614 |  |
 470 - 690 ms | 580 ms | 0.0001 | 1.4888 |  | | | | 880 - 990 ms | 880 ms | 0.015 | -0.6665 |  | | | ||  | **posterior** | 190 - 690 ms | 580 ms | 0.0002 | 1.6366 | 200 - 330 ms | 250 ms | 0.009 | 1.2744 |
 720 - 780 ms | 740 ms | 0.0314 | 1.0619 | 350 - 610 ms | 480 ms | 0.0001 | 1.935 | 820 - 890 ms | 840 ms | 0.048 | 0.8134 |  | | | |

  

E) **familiar vs. remembered objects**

|  | time window | peak latency | cluster *p* | peak Cohen's *d* |  | | | |  |
| **all electrodes** | 190 - 410 ms | 300 ms | 0.0021 | 0.9267 |  | | | |  |
 660 - 840 ms | 740 ms | 0.0034 | -0.5453 |  | | | ||  |  | | | | | | | | |
|  | **left hemisphere** | | | | **right hemisphere** | | | |  |
|  | time window | peak latency | cluster *p* | peak Cohen's *d* | time window | peak latency | cluster *p* | peak Cohen's *d* |  |
| **anterior** | 430 - 880 ms | 880 ms | 0.0002 | -0.4677 | 220 - 350 ms | 260 ms | 0.0113 | 1.034 |  |
  | | | | 520 - 900 ms | 900 ms | 0.0001 | -0.5649 ||  | **central** | 200 - 490 ms | 390 ms | 0.0001 | 1.6872 | 210 - 440 ms | 300 ms | 0.0014 | 1.1087 |
 550 - 720 ms | 720 ms | 0.0024 | -0.4914 | 540 - 840 ms | 840 ms | 0.0001 | -0.5128 || **posterior** |  | 280 - 430 ms | 300 ms | 0.0054 | 1.0042 | 240 - 410 ms | 380 ms | 0.0021 | 1.4639 |
 670 - 790 ms | 740 ms | 0.0209 | -0.4692 | 550 - 640 ms | 550 ms | 0.0308 | -0.4951 |  | | | | 660 - 910 ms | 910 ms | 0.0019 | -0.4749 |

  
