## Supplementary Table for "Shared neural codes of recognition memory": Supplementary_Table_3.html

  


### Supplementary Table 3.

  

Results of the main temporal generalization analyses  

#### A) personally familiar and unfamiliar faces (leave-one-subject-out)

  

  

left anterior

|  | train times | test times | cluster p | cluster p-star | peak Cohen's d | direction |
| --- | --- | --- | --- | --- | --- | --- |
| #1 | 190 to 990 ms | 200 to 990 ms | 0.0001 | \*\*\* | 1.575 | positive |

  

right anterior

|  | train times | test times | cluster p | cluster p-star | peak Cohen's d | direction |
| --- | --- | --- | --- | --- | --- | --- |
| #1 | -80 to 130 ms | 340 to 990 ms | 0.0434 | \* | 1.3928 | positive |
| #2 | 180 to 990 ms | 190 to 990 ms | 0.0001 | \*\*\* | 1.7011 | positive |

  

left central

|  | train times | test times | cluster p | cluster p-star | peak Cohen's d | direction |
| --- | --- | --- | --- | --- | --- | --- |
| #1 | 160 to 990 ms | 200 to 990 ms | 0.0001 | \*\*\* | 2.287 | positive |

  

right central

|  | train times | test times | cluster p | cluster p-star | peak Cohen's d | direction |
| --- | --- | --- | --- | --- | --- | --- |
| #1 | 170 to 990 ms | 200 to 990 ms | 0.0001 | \*\*\* | 2.3225 | positive |

  

left posterior

|  | train times | test times | cluster p | cluster p-star | peak Cohen's d | direction |
| --- | --- | --- | --- | --- | --- | --- |
| #1 | 180 to 990 ms | 190 to 990 ms | 0.0001 | \*\*\* | 1.9169 | positive |

  

right posterior

|  | train times | test times | cluster p | cluster p-star | peak Cohen's d | direction |
| --- | --- | --- | --- | --- | --- | --- |
| #1 | 180 to 990 ms | 200 to 990 ms | 0.0001 | \*\*\* | 1.9952 | positive |

  

all electrodes

|  | train times | test times | cluster p | cluster p-star | peak Cohen's d | direction |
| --- | --- | --- | --- | --- | --- | --- |
| #1 | 170 to 990 ms | 200 to 990 ms | 0.0001 | \*\*\* | 2.5397 | positive |

#### B) trained on experimentally familiarized face and unfamiliar faces, tested on personally familiar and unfamiliar faces

  

  

left anterior

|  | train times | test times | cluster p | cluster p-star | peak Cohen's d | direction |
| --- | --- | --- | --- | --- | --- | --- |
| #1 | 160 to 360 ms | 230 to 990 ms | 0.0192 | \* | 1.1556 | positive |
| #2 | 390 to 630 ms | 310 to 990 ms | 0.0086 | \*\* | 1.3668 | positive |
| #3 | 650 to 1290 ms | 210 to 990 ms | 0.0001 | \*\*\* | -2.0133 | negative |

  

right anterior

|  | train times | test times | cluster p | cluster p-star | peak Cohen's d | direction |
| --- | --- | --- | --- | --- | --- | --- |
| #1 | 400 to 680 ms | 190 to 990 ms | 0.0065 | \*\* | 1.5612 | positive |
| #2 | 720 to 1290 ms | 190 to 990 ms | 0.0001 | \*\*\* | -1.9561 | negative |

  

left central

|  | train times | test times | cluster p | cluster p-star | peak Cohen's d | direction |
| --- | --- | --- | --- | --- | --- | --- |
| #2 | 390 to 790 ms | 200 to 990 ms | 0.0001 | \*\*\* | 2.2455 | positive |
| #3 | -40 to 180 ms | 200 to 990 ms | 0.0366 | \* | -1.5342 | negative |
| #1 | 200 to 330 ms | 210 to 990 ms | 0.0337 | \* | 1.3869 | positive |
| #4 | 800 to 1290 ms | 200 to 990 ms | 0.0001 | \*\*\* | -1.6363 | negative |

  

right central

|  | train times | test times | cluster p | cluster p-star | peak Cohen's d | direction |
| --- | --- | --- | --- | --- | --- | --- |
| #1 | 130 to 330 ms | 200 to 990 ms | 0.0158 | \* | 1.8684 | positive |
| #2 | 380 to 840 ms | 200 to 990 ms | 0.0001 | \*\*\* | 2.1075 | positive |
| #3 | 800 to 1290 ms | 210 to 990 ms | 0.0001 | \*\*\* | -2.481 | negative |

  

left posterior

|  | train times | test times | cluster p | cluster p-star | peak Cohen's d | direction |
| --- | --- | --- | --- | --- | --- | --- |
| #1 | 90 to 340 ms | 160 to 990 ms | 0.0447 | \* | 0.9889 | positive |
| #2 | 370 to 790 ms | 150 to 990 ms | 0.0001 | \*\*\* | 1.7512 | positive |
| #3 | 750 to 1290 ms | 210 to 990 ms | 0.0017 | \*\* | -1.5627 | negative |

  

right posterior

|  | train times | test times | cluster p | cluster p-star | peak Cohen's d | direction |
| --- | --- | --- | --- | --- | --- | --- |
| #1 | 90 to 840 ms | 200 to 990 ms | 0.0001 | \*\*\* | 2.5373 | positive |
| #2 | 850 to 1290 ms | 200 to 990 ms | 0.002 | \*\* | -2.0752 | negative |

  

all electrodes

|  | train times | test times | cluster p | cluster p-star | peak Cohen's d | direction |
| --- | --- | --- | --- | --- | --- | --- |
| #1 | 90 to 830 ms | 190 to 990 ms | 0.0001 | \*\*\* | 1.9626 | positive |
| #2 | 820 to 1290 ms | 220 to 990 ms | 0.014 | \* | -1.392 | negative |

#### C) trained on famous and unfamiliar faces, tested on personally familiar and unfamiliar faces

  

  

left anterior

|  | train times | test times | cluster p | cluster p-star | peak Cohen's d | direction |
| --- | --- | --- | --- | --- | --- | --- |
| #1 | 130 to 1290 ms | 200 to 990 ms | 0.0001 | \*\*\* | 1.6241 | positive |

  

right anterior

|  | train times | test times | cluster p | cluster p-star | peak Cohen's d | direction |
| --- | --- | --- | --- | --- | --- | --- |
| #1 | 210 to 1290 ms | 200 to 990 ms | 0.0001 | \*\*\* | 1.5742 | positive |

  

left central

|  | train times | test times | cluster p | cluster p-star | peak Cohen's d | direction |
| --- | --- | --- | --- | --- | --- | --- |
| #1 | 190 to 1290 ms | 200 to 990 ms | 0.0001 | \*\*\* | 1.9269 | positive |

  

right central

|  | train times | test times | cluster p | cluster p-star | peak Cohen's d | direction |
| --- | --- | --- | --- | --- | --- | --- |
| #1 | 200 to 1290 ms | 200 to 990 ms | 0.0001 | \*\*\* | 2.3178 | positive |

  

left posterior

|  | train times | test times | cluster p | cluster p-star | peak Cohen's d | direction |
| --- | --- | --- | --- | --- | --- | --- |
| #1 | 190 to 1290 ms | 190 to 990 ms | 0.0002 | \*\*\* | 1.4136 | positive |

  

right posterior

|  | train times | test times | cluster p | cluster p-star | peak Cohen's d | direction |
| --- | --- | --- | --- | --- | --- | --- |
| #1 | 200 to 1290 ms | 190 to 990 ms | 0.0001 | \*\*\* | 1.7371 | positive |

  

all electrodes

|  | train times | test times | cluster p | cluster p-star | peak Cohen's d | direction |
| --- | --- | --- | --- | --- | --- | --- |
| #1 | 200 to 1290 ms | 200 to 990 ms | 0.0002 | \*\*\* | 1.5614 | positive |

#### D) trained on familiar and unfamiliar music, tested on personally familiar and unfamiliar faces

  

  

left anterior

|  | train times | test times | cluster p | cluster p-star | peak Cohen's d | direction |
| --- | --- | --- | --- | --- | --- | --- |
| #3 | 1210 to 1490 ms | 210 to 650 ms | 0.0298 | \* | 0.9669 | positive |
| #4 | 1250 to 1420 ms | 320 to 990 ms | 0.02 | \* | 1.1239 | positive |
| #6 | 170 to 250 ms | 210 to 990 ms | 0.0133 | \* | -1.2316 | negative |
| #7 | 970 to 1050 ms | 350 to 930 ms | 0.0258 | \* | -1.8357 | negative |
| #1 | 290 to 340 ms | 200 to 990 ms | 0.0409 | \* | 1.3809 | positive |
| #2 | 680 to 870 ms | 200 to 990 ms | 0.0079 | \*\* | 1.294 | positive |
| #5 | -60 to 40 ms | 210 to 990 ms | 0.0198 | \* | -1.4441 | negative |

  

right anterior

|  | train times | test times | cluster p | cluster p-star | peak Cohen's d | direction |
| --- | --- | --- | --- | --- | --- | --- |
| #4 | 1210 to 1490 ms | 210 to 990 ms | 0.0008 | \*\*\* | 1.2164 | positive |
| #5 | -170 to -100 ms | 200 to 880 ms | 0.0492 | \* | -1.1801 | negative |
| #6 | -20 to 30 ms | 320 to 940 ms | 0.0464 | \* | -1.5606 | negative |
| #7 | 60 to 110 ms | 290 to 930 ms | 0.0414 | \* | -1.5037 | negative |
| #1 | 290 to 330 ms | 190 to 990 ms | 0.0284 | \* | 1.6369 | positive |
| #2 | 360 to 850 ms | 190 to 990 ms | 0.0001 | \*\*\* | 1.701 | positive |
| #3 | 980 to 1090 ms | 200 to 990 ms | 0.0056 | \*\* | 1.4868 | positive |

  

left central

|  | train times | test times | cluster p | cluster p-star | peak Cohen's d | direction |
| --- | --- | --- | --- | --- | --- | --- |
| #1 | -170 to -90 ms | 220 to 990 ms | 0.0021 | \*\* | 2.1057 | positive |
| #3 | 110 to 160 ms | 310 to 870 ms | 0.0087 | \*\* | 1.6824 | positive |
| #4 | 290 to 390 ms | 200 to 990 ms | 0.0005 | \*\*\* | 1.7235 | positive |
| #6 | 530 to 560 ms | 210 to 980 ms | 0.021 | \* | 1.7006 | positive |
| #7 | 610 to 670 ms | 180 to 990 ms | 0.0081 | \*\* | 1.8175 | positive |
| #8 | 690 to 780 ms | 200 to 990 ms | 0.0001 | \*\*\* | 2.7435 | positive |
| #10 | 1150 to 1280 ms | 190 to 990 ms | 0.0005 | \*\*\* | 1.1202 | positive |
| #11 | 1310 to 1340 ms | 220 to 990 ms | 0.0136 | \* | 1.4993 | positive |
| #13 | -70 to -20 ms | 210 to 990 ms | 0.0021 | \*\* | -1.6134 | negative |
| #15 | 570 to 610 ms | 400 to 760 ms | 0.035 | \* | -1.1723 | negative |
| #16 | 790 to 810 ms | 350 to 980 ms | 0.0335 | \* | -1.4933 | negative |
| #17 | 1040 to 1090 ms | 200 to 990 ms | 0.0078 | \*\* | -2.4039 | negative |
| #2 | -10 to 20 ms | 200 to 990 ms | 0.0059 | \*\* | 2.2444 | positive |
| #5 | 420 to 470 ms | 290 to 820 ms | 0.0144 | \* | 1.3923 | positive |
| #9 | 820 to 910 ms | 190 to 990 ms | 0.0019 | \*\* | 1.6544 | positive |
| #12 | 1360 to 1460 ms | 240 to 990 ms | 0.0017 | \*\* | 1.7164 | positive |
| #14 | 260 to 290 ms | 300 to 770 ms | 0.0335 | \* | -1.2415 | negative |

  

right central

|  | train times | test times | cluster p | cluster p-star | peak Cohen's d | direction |
| --- | --- | --- | --- | --- | --- | --- |
| #3 | 1140 to 1490 ms | 180 to 990 ms | 0.0021 | \*\* | 2.0276 | positive |
| #4 | -90 to -40 ms | 190 to 990 ms | 0.0462 | \* | -1.9109 | negative |
| #5 | -10 to 50 ms | 300 to 990 ms | 0.0481 | \* | -1.6304 | negative |
| #1 | 50 to 180 ms | 210 to 990 ms | 0.0304 | \* | 2.5332 | positive |
| #2 | 290 to 1120 ms | 190 to 990 ms | 0.0001 | \*\*\* | 2.0454 | positive |

  

left posterior

|  | train times | test times | cluster p | cluster p-star | peak Cohen's d | direction |
| --- | --- | --- | --- | --- | --- | --- |
| #1 | -150 to -90 ms | 180 to 990 ms | 0.0324 | \* | 1.334 | positive |
| #2 | 120 to 160 ms | 170 to 990 ms | 0.0338 | \* | 1.6849 | positive |
| #3 | 200 to 240 ms | 290 to 990 ms | 0.0469 | \* | 1.7549 | positive |
| #4 | 290 to 390 ms | 200 to 990 ms | 0.0149 | \* | 1.8097 | positive |
| #5 | 420 to 500 ms | 290 to 990 ms | 0.0215 | \* | 1.597 | positive |
| #6 | 520 to 790 ms | 290 to 990 ms | 0.0056 | \*\* | 1.4373 | positive |
| #7 | 820 to 870 ms | 280 to 990 ms | 0.0314 | \* | 1.6256 | positive |
| #8 | 900 to 1430 ms | 180 to 990 ms | 0.0001 | \*\*\* | 1.8287 | positive |
| #9 | 1470 to 1490 ms | 190 to 990 ms | 0.0366 | \* | 1.9489 | positive |

  

right posterior

|  | train times | test times | cluster p | cluster p-star | peak Cohen's d | direction |
| --- | --- | --- | --- | --- | --- | --- |
| #2 | 110 to 160 ms | 210 to 990 ms | 0.0314 | \* | 1.4044 | positive |
| #5 | 510 to 850 ms | 200 to 990 ms | 0.0008 | \*\*\* | 2.0669 | positive |
| #6 | 900 to 970 ms | 210 to 990 ms | 0.0378 | \* | 1.5154 | positive |
| #7 | 990 to 1050 ms | 200 to 990 ms | 0.0251 | \* | 2.1035 | positive |
| #8 | 1050 to 1110 ms | 190 to 990 ms | 0.0278 | \* | 1.633 | positive |
| #9 | 1130 to 1230 ms | 200 to 990 ms | 0.0275 | \* | 1.7788 | positive |
| #10 | 1290 to 1340 ms | 200 to 990 ms | 0.0268 | \* | 1.9658 | positive |
| #11 | 1400 to 1490 ms | 190 to 990 ms | 0.0121 | \* | 1.7807 | positive |
| #12 | -200 to -140 ms | 190 to 990 ms | 0.0321 | \* | -1.2123 | negative |
| #13 | -80 to -20 ms | 290 to 990 ms | 0.0436 | \* | -1.0437 | negative |
| #14 | 40 to 100 ms | 220 to 990 ms | 0.0387 | \* | -1.2114 | negative |
| #1 | -130 to -90 ms | 210 to 990 ms | 0.0488 | \* | 1.2719 | positive |
| #3 | 240 to 280 ms | 200 to 990 ms | 0.0415 | \* | 2.0304 | positive |
| #4 | 300 to 490 ms | 200 to 990 ms | 0.0037 | \*\* | 2.0049 | positive |

  

all electrodes

|  | train times | test times | cluster p | cluster p-star | peak Cohen's d | direction |
| --- | --- | --- | --- | --- | --- | --- |
| #2 | 290 to 1100 ms | 200 to 990 ms | 0.0001 | \*\*\* | 2.332 | positive |
| #4 | -200 to -140 ms | 200 to 990 ms | 0.0293 | \* | -2.0732 | negative |
| #5 | 20 to 190 ms | 200 to 990 ms | 0.0259 | \* | -1.7707 | negative |
| #1 | -140 to -70 ms | 190 to 990 ms | 0.0264 | \* | 1.706 | positive |
| #3 | 1130 to 1490 ms | 190 to 990 ms | 0.0029 | \*\* | 2.0971 | positive |

#### E) trained on remembered and forgotten object-scene associations, tested on personally familiar and unfamiliar faces

  

  

left anterior

|  | train times | test times | cluster p | cluster p-star | peak Cohen's d | direction |
| --- | --- | --- | --- | --- | --- | --- |
| #1 | 250 to 710 ms | 310 to 990 ms | 0.0008 | \*\*\* | 1.6546 | positive |
| #2 | 960 to 1280 ms | 320 to 650 ms | 0.0199 | \* | 1.3652 | positive |

  

right anterior

|  | train times | test times | cluster p | cluster p-star | peak Cohen's d | direction |
| --- | --- | --- | --- | --- | --- | --- |
| #1 | 320 to 1290 ms | 220 to 990 ms | 0.0004 | \*\*\* | 1.3105 | positive |

  

left central

|  | train times | test times | cluster p | cluster p-star | peak Cohen's d | direction |
| --- | --- | --- | --- | --- | --- | --- |
| #1 | -30 to 200 ms | 200 to 990 ms | 0.01 | \* | 2.1018 | positive |
| #2 | 230 to 700 ms | 200 to 990 ms | 0.0001 | \*\*\* | 2.0132 | positive |
| #3 | 810 to 1290 ms | 200 to 990 ms | 0.0001 | \*\*\* | 1.9923 | positive |

  

right central

|  | train times | test times | cluster p | cluster p-star | peak Cohen's d | direction |
| --- | --- | --- | --- | --- | --- | --- |
| #1 | -200 to -110 ms | 200 to 990 ms | 0.0437 | \* | 2.0496 | positive |
| #2 | 0 to 200 ms | 200 to 990 ms | 0.0061 | \*\* | 1.9329 | positive |
| #3 | 330 to 720 ms | 200 to 990 ms | 0.0001 | \*\*\* | 2.2679 | positive |
| #4 | 810 to 900 ms | 200 to 990 ms | 0.0228 | \* | 1.9434 | positive |
| #5 | 930 to 1290 ms | 200 to 990 ms | 0.0001 | \*\*\* | 1.9585 | positive |

  

left posterior

|  | train times | test times | cluster p | cluster p-star | peak Cohen's d | direction |
| --- | --- | --- | --- | --- | --- | --- |
| #1 | 90 to 200 ms | 170 to 990 ms | 0.017 | \* | 1.6575 | positive |
| #2 | 230 to 670 ms | 180 to 990 ms | 0.0003 | \*\*\* | 1.5454 | positive |
| #3 | 760 to 1130 ms | 190 to 670 ms | 0.0214 | \* | -1.0434 | negative |

  

right posterior

|  | train times | test times | cluster p | cluster p-star | peak Cohen's d | direction |
| --- | --- | --- | --- | --- | --- | --- |
| #1 | 30 to 210 ms | 210 to 990 ms | 0.0221 | \* | 1.8439 | positive |
| #2 | 230 to 710 ms | 200 to 990 ms | 0.0002 | \*\*\* | 2.0679 | positive |
| #3 | 1040 to 1290 ms | 200 to 990 ms | 0.0008 | \*\*\* | 1.8783 | positive |

  

all electrodes

|  | train times | test times | cluster p | cluster p-star | peak Cohen's d | direction |
| --- | --- | --- | --- | --- | --- | --- |
| #1 | -20 to 210 ms | 300 to 990 ms | 0.0056 | \*\* | 1.5089 | positive |
| #2 | 310 to 640 ms | 230 to 990 ms | 0.0005 | \*\*\* | 1.9731 | positive |
| #4 | 650 to 1030 ms | 200 to 710 ms | 0.0425 | \* | -1.3005 | negative |
| #3 | 1130 to 1290 ms | 290 to 780 ms | 0.0249 | \* | 1.3681 | positive |

#### F) trained on familiar and novel objects, tested on personally familiar and unfamiliar faces

  

  

left anterior

|  | train times | test times | cluster p | cluster p-star | peak Cohen's d | direction |
| --- | --- | --- | --- | --- | --- | --- |
| #1 | 180 to 330 ms | 200 to 940 ms | 0.0339 | \* | 1.5279 | positive |
| #2 | 380 to 570 ms | 300 to 950 ms | 0.041 | \* | 1.5298 | positive |
| #3 | 580 to 1100 ms | 330 to 870 ms | 0.0032 | \*\* | -1.1905 | negative |

  

right anterior

|  | train times | test times | cluster p | cluster p-star | peak Cohen's d | direction |
| --- | --- | --- | --- | --- | --- | --- |
| #2 | -70 to 100 ms | 190 to 990 ms | 0.0306 | \* | -1.1932 | negative |
| #1 | 160 to 620 ms | 200 to 990 ms | 0.0001 | \*\*\* | 1.7042 | positive |
| #3 | 620 to 1260 ms | 310 to 960 ms | 0.0026 | \*\* | -1.0951 | negative |

  

left central

|  | train times | test times | cluster p | cluster p-star | peak Cohen's d | direction |
| --- | --- | --- | --- | --- | --- | --- |
| #1 | 100 to 700 ms | 200 to 990 ms | 0.0001 | \*\*\* | 1.9715 | positive |
| #2 | 670 to 1250 ms | 200 to 990 ms | 0.0001 | \*\*\* | -1.7875 | negative |

  

right central

|  | train times | test times | cluster p | cluster p-star | peak Cohen's d | direction |
| --- | --- | --- | --- | --- | --- | --- |
| #1 | 90 to 670 ms | 190 to 990 ms | 0.0001 | \*\*\* | 3.0205 | positive |
| #2 | 660 to 1270 ms | 190 to 990 ms | 0.0001 | \*\*\* | -1.9195 | negative |

  

left posterior

|  | train times | test times | cluster p | cluster p-star | peak Cohen's d | direction |
| --- | --- | --- | --- | --- | --- | --- |
| #1 | 160 to 630 ms | 180 to 990 ms | 0.0026 | \*\* | 1.6479 | positive |
| #2 | 650 to 1290 ms | 210 to 990 ms | 0.0001 | \*\*\* | -1.4657 | negative |

  

right posterior

|  | train times | test times | cluster p | cluster p-star | peak Cohen's d | direction |
| --- | --- | --- | --- | --- | --- | --- |
| #1 | 170 to 670 ms | 200 to 990 ms | 0.0001 | \*\*\* | 2.0392 | positive |
| #2 | 730 to 1290 ms | 200 to 990 ms | 0.0001 | \*\*\* | -1.7523 | negative |

  

all electrodes

|  | train times | test times | cluster p | cluster p-star | peak Cohen's d | direction |
| --- | --- | --- | --- | --- | --- | --- |
| #1 | 130 to 770 ms | 160 to 990 ms | 0.0001 | \*\*\* | 1.6322 | positive |
| #2 | 850 to 1290 ms | 240 to 990 ms | 0.0014 | \*\* | -1.5874 | negative |

#### G) trained on remembered and novel objects, tested on personally familiar and unfamiliar faces

  

  

left anterior

|  | train times | test times | cluster p | cluster p-star | peak Cohen's d | direction |
| --- | --- | --- | --- | --- | --- | --- |
| #1 | 360 to 1130 ms | 220 to 990 ms | 0.0002 | \*\*\* | 1.413 | positive |

  

right anterior

|  | train times | test times | cluster p | cluster p-star | peak Cohen's d | direction |
| --- | --- | --- | --- | --- | --- | --- |
| #2 | -70 to 40 ms | 210 to 990 ms | 0.0483 | \* | -1.5318 | negative |
| #1 | 410 to 1030 ms | 210 to 990 ms | 0.0001 | \*\*\* | 1.6908 | positive |

  

left central

|  | train times | test times | cluster p | cluster p-star | peak Cohen's d | direction |
| --- | --- | --- | --- | --- | --- | --- |
| #2 | 70 to 310 ms | 190 to 990 ms | 0.013 | \* | -2.2481 | negative |
| #1 | 400 to 710 ms | 200 to 990 ms | 0.0013 | \*\* | 2.1479 | positive |
| #3 | 750 to 1290 ms | 200 to 990 ms | 0.0001 | \*\*\* | -1.669 | negative |

  

right central

|  | train times | test times | cluster p | cluster p-star | peak Cohen's d | direction |
| --- | --- | --- | --- | --- | --- | --- |
| #2 | 50 to 390 ms | 190 to 990 ms | 0.0127 | \* | -1.6751 | negative |
| #1 | 400 to 760 ms | 200 to 990 ms | 0.0003 | \*\*\* | 1.8793 | positive |
| #3 | 770 to 1290 ms | 190 to 990 ms | 0.0001 | \*\*\* | -1.9294 | negative |

  

left posterior

|  | train times | test times | cluster p | cluster p-star | peak Cohen's d | direction |
| --- | --- | --- | --- | --- | --- | --- |
| #1 | 270 to 680 ms | 210 to 990 ms | 0.0168 | \* | 1.7546 | positive |
| #2 | 690 to 1290 ms | 190 to 990 ms | 0.0001 | \*\*\* | -1.5542 | negative |

  

right posterior

|  | train times | test times | cluster p | cluster p-star | peak Cohen's d | direction |
| --- | --- | --- | --- | --- | --- | --- |
| #1 | 370 to 780 ms | 180 to 990 ms | 0.0002 | \*\*\* | 2.1264 | positive |
| #2 | 770 to 1290 ms | 200 to 990 ms | 0.0001 | \*\*\* | -1.7566 | negative |

  

all electrodes

|  | train times | test times | cluster p | cluster p-star | peak Cohen's d | direction |
| --- | --- | --- | --- | --- | --- | --- |
| #1 | 400 to 760 ms | 280 to 990 ms | 0.0046 | \*\* | 1.7849 | positive |
| #2 | 760 to 1290 ms | 180 to 990 ms | 0.0002 | \*\*\* | -1.6403 | negative |
