## Supplementary Table for "Shared neural codes of recognition memory": Supplementary_Table_4.html

  


### Supplementary Table 4.

  

Results of the temporal generalization analyses, trained on the familiar - remembered object dataset  

#### A) trained on FA familiar and novel objects, tested on personally familiar and unfamiliar faces

  

  

left anterior

|  | train times | test times | cluster p | cluster p-star | peak Cohen's d | direction |
| --- | --- | --- | --- | --- | --- | --- |
| #1 | 140 to 630 ms | 190 to 990 ms | 0.0011 | \*\* | 1.2923 | positive |
| #2 | 1070 to 1290 ms | 210 to 990 ms | 0.0148 | \* | 1.0452 | positive |

  

right anterior

|  | train times | test times | cluster p | cluster p-star | peak Cohen's d | direction |
| --- | --- | --- | --- | --- | --- | --- |
| #2 | 1100 to 1290 ms | 200 to 990 ms | 0.0221 | \* | 1.8181 | positive |
| #3 | -60 to 70 ms | 210 to 990 ms | 0.041 | \* | -1.3814 | negative |
| #1 | 170 to 690 ms | 210 to 990 ms | 0.0016 | \*\* | 1.229 | positive |

  

left central

|  | train times | test times | cluster p | cluster p-star | peak Cohen's d | direction |
| --- | --- | --- | --- | --- | --- | --- |
| #1 | 70 to 1070 ms | 190 to 990 ms | 0.0001 | \*\*\* | 2.1269 | positive |
| #2 | 1110 to 1290 ms | 200 to 990 ms | 0.0152 | \* | 2.0557 | positive |

  

right central

|  | train times | test times | cluster p | cluster p-star | peak Cohen's d | direction |
| --- | --- | --- | --- | --- | --- | --- |
| #2 | -40 to 60 ms | 200 to 990 ms | 0.0479 | \* | -1.3535 | negative |
| #1 | 140 to 950 ms | 190 to 990 ms | 0.0001 | \*\*\* | 2.8085 | positive |

  

left posterior

|  | train times | test times | cluster p | cluster p-star | peak Cohen's d | direction |
| --- | --- | --- | --- | --- | --- | --- |
| #1 | 130 to 860 ms | 150 to 990 ms | 0.0001 | \*\*\* | 1.9133 | positive |
| #2 | 1080 to 1290 ms | 180 to 990 ms | 0.024 | \* | 1.5747 | positive |

  

right posterior

|  | train times | test times | cluster p | cluster p-star | peak Cohen's d | direction |
| --- | --- | --- | --- | --- | --- | --- |
| #1 | 60 to 860 ms | 200 to 990 ms | 0.0003 | \*\*\* | 1.3801 | positive |
| #2 | 1130 to 1290 ms | 210 to 990 ms | 0.0404 | \* | 1.8095 | positive |

  

all electrodes

|  | train times | test times | cluster p | cluster p-star | peak Cohen's d | direction |
| --- | --- | --- | --- | --- | --- | --- |
| #1 | 50 to 130 ms | 220 to 990 ms | 0.0455 | \* | 1.7753 | positive |
| #2 | 150 to 950 ms | 170 to 990 ms | 0.0002 | \*\*\* | 1.5126 | positive |
| #3 | 1120 to 1290 ms | 180 to 990 ms | 0.0074 | \*\* | 1.834 | positive |

#### B) trained on FA remembered and novel objects, tested on personally familiar and unfamiliar faces

  

  

left anterior

|  | train times | test times | cluster p | cluster p-star | peak Cohen's d | direction |
| --- | --- | --- | --- | --- | --- | --- |
| #2 | -10 to 170 ms | 220 to 990 ms | 0.0044 | \*\* | -1.5554 | negative |
| #1 | 530 to 880 ms | 200 to 630 ms | 0.0046 | \*\* | 1.2037 | positive |

  

right anterior

|  | train times | test times | cluster p | cluster p-star | peak Cohen's d | direction |
| --- | --- | --- | --- | --- | --- | --- |
| #2 | 460 to 1290 ms | 180 to 990 ms | 0.0001 | \*\*\* | 1.817 | positive |
| #3 | -40 to 100 ms | 160 to 990 ms | 0.0411 | \* | -1.5011 | negative |
| #1 | 170 to 330 ms | 190 to 990 ms | 0.0397 | \* | 1.3271 | positive |

  

left central

|  | train times | test times | cluster p | cluster p-star | peak Cohen's d | direction |
| --- | --- | --- | --- | --- | --- | --- |
| #1 | -80 to 20 ms | 200 to 990 ms | 0.0327 | \* | 1.9436 | positive |
| #2 | 150 to 1020 ms | 190 to 990 ms | 0.0001 | \*\*\* | 2.2087 | positive |
| #3 | 1190 to 1290 ms | 360 to 960 ms | 0.0469 | \* | -1.4507 | negative |

  

right central

|  | train times | test times | cluster p | cluster p-star | peak Cohen's d | direction |
| --- | --- | --- | --- | --- | --- | --- |
| #1 | 160 to 1020 ms | 180 to 990 ms | 0.0001 | \*\*\* | 2.216 | positive |

  

left posterior

|  | train times | test times | cluster p | cluster p-star | peak Cohen's d | direction |
| --- | --- | --- | --- | --- | --- | --- |
| #1 | -70 to 0 ms | 220 to 990 ms | 0.049 | \* | 0.7593 | positive |
| #2 | 50 to 90 ms | 180 to 990 ms | 0.025 | \* | 1.9388 | positive |
| #4 | 620 to 810 ms | 200 to 990 ms | 0.0009 | \*\*\* | 1.8038 | positive |
| #5 | 110 to 160 ms | 280 to 990 ms | 0.0304 | \* | -1.3438 | negative |
| #3 | 490 to 600 ms | 220 to 990 ms | 0.0105 | \* | 1.3569 | positive |
| #6 | 1110 to 1290 ms | 240 to 990 ms | 0.0082 | \*\* | -1.5147 | negative |

  

right posterior

|  | train times | test times | cluster p | cluster p-star | peak Cohen's d | direction |
| --- | --- | --- | --- | --- | --- | --- |
| #1 | 170 to 880 ms | 200 to 990 ms | 0.0001 | \*\*\* | 2.523 | positive |

  

all electrodes

|  | train times | test times | cluster p | cluster p-star | peak Cohen's d | direction |
| --- | --- | --- | --- | --- | --- | --- |
| #3 | 100 to 150 ms | 210 to 990 ms | 0.0399 | \* | -1.5818 | negative |
| #1 | 170 to 1100 ms | 190 to 990 ms | 0.0001 | \*\*\* | 2.3532 | positive |
| #2 | -10 to 60 ms | 210 to 980 ms | 0.0443 | \* | -0.9062 | negative |

#### C) trained on forgotten vs. novel objects, tested on personally familiar and unfamiliar faces

  

  

left anterior

|  | train times | test times | cluster p | cluster p-star | peak Cohen's d | direction |
| --- | --- | --- | --- | --- | --- | --- |
| #1 | -70 to 100 ms | 220 to 870 ms | 0.045 | \* | -1.2241 | negative |
| #2 | 1030 to 1200 ms | 320 to 990 ms | 0.034 | \* | -1.808 | negative |

  

right anterior

|  | train times | test times | cluster p | cluster p-star | peak Cohen's d | direction |
| --- | --- | --- | --- | --- | --- | --- |
| #1 | 450 to 600 ms | 360 to 990 ms | 0.0491 | \* | 0.9455 | positive |

  

left central

|  | train times | test times | cluster p | cluster p-star | peak Cohen's d | direction |
| --- | --- | --- | --- | --- | --- | --- |
| #1 | 180 to 870 ms | 190 to 990 ms | 0.0001 | \*\*\* | 2.2515 | positive |

  

right central

|  | train times | test times | cluster p | cluster p-star | peak Cohen's d | direction |
| --- | --- | --- | --- | --- | --- | --- |
| #1 | 160 to 240 ms | 210 to 990 ms | 0.0452 | \* | 1.6701 | positive |
| #2 | 270 to 780 ms | 190 to 990 ms | 0.0001 | \*\*\* | 2.2984 | positive |

  

left posterior

|  | train times | test times | cluster p | cluster p-star | peak Cohen's d | direction |
| --- | --- | --- | --- | --- | --- | --- |
| #1 | 270 to 690 ms | 180 to 990 ms | 0.0001 | \*\*\* | 1.5974 | positive |
| #2 | 840 to 1010 ms | 220 to 990 ms | 0.0363 | \* | -1.2249 | negative |

  

right posterior

|  | train times | test times | cluster p | cluster p-star | peak Cohen's d | direction |
| --- | --- | --- | --- | --- | --- | --- |
| #1 | 180 to 850 ms | 190 to 990 ms | 0.0001 | \*\*\* | 2.2611 | positive |
| #2 | 1030 to 1190 ms | 220 to 990 ms | 0.0376 | \* | -0.786 | negative |

  

all electrodes

|  | train times | test times | cluster p | cluster p-star | peak Cohen's d | direction |
| --- | --- | --- | --- | --- | --- | --- |
| #1 | 260 to 860 ms | 190 to 990 ms | 0.0038 | \*\* | 1.0429 | positive |

#### D) trained on objects, incorrect answers with subjective labels, tested on personally familiar and unfamiliar faces

  

  

left anterior

|  | train times | test times | cluster p | cluster p-star | peak Cohen's d | direction |
| --- | --- | --- | --- | --- | --- | --- |
| #1 | 160 to 280 ms | 220 to 990 ms | 0.0182 | \* | 1.1525 | positive |
| #2 | 600 to 1140 ms | 200 to 680 ms | 0.0129 | \* | -1.3456 | negative |

  

right anterior

|  | train times | test times | cluster p | cluster p-star | peak Cohen's d | direction |
| --- | --- | --- | --- | --- | --- | --- |
| #2 | 370 to 640 ms | 280 to 950 ms | 0.0032 | \*\* | 1.5441 | positive |
| #3 | -200 to -140 ms | 220 to 990 ms | 0.0395 | \* | -1.0576 | negative |
| #1 | 140 to 280 ms | 210 to 990 ms | 0.0063 | \*\* | 1.2441 | positive |
| #4 | 610 to 1080 ms | 200 to 990 ms | 0.0016 | \*\* | -1.2422 | negative |

  

left central

|  | train times | test times | cluster p | cluster p-star | peak Cohen's d | direction |
| --- | --- | --- | --- | --- | --- | --- |
| #1 | 70 to 450 ms | 200 to 780 ms | 0.0031 | \*\* | 1.6184 | positive |
| #2 | 460 to 870 ms | 210 to 990 ms | 0.0015 | \*\* | 1.9024 | positive |
| #3 | 870 to 1020 ms | 300 to 990 ms | 0.0145 | \* | -1.6262 | negative |

  

right central

|  | train times | test times | cluster p | cluster p-star | peak Cohen's d | direction |
| --- | --- | --- | --- | --- | --- | --- |
| #1 | -170 to -100 ms | 200 to 990 ms | 0.0471 | \* | 1.824 | positive |
| #2 | 70 to 310 ms | 190 to 990 ms | 0.0113 | \* | 2.418 | positive |
| #3 | 340 to 890 ms | 200 to 990 ms | 0.0001 | \*\*\* | 2.2634 | positive |
| #4 | 900 to 1000 ms | 300 to 990 ms | 0.0458 | \* | -1.3571 | negative |

  

left posterior

|  | train times | test times | cluster p | cluster p-star | peak Cohen's d | direction |
| --- | --- | --- | --- | --- | --- | --- |
| #1 | -70 to 800 ms | 150 to 990 ms | 0.0001 | \*\*\* | 1.8526 | positive |

  

right posterior

|  | train times | test times | cluster p | cluster p-star | peak Cohen's d | direction |
| --- | --- | --- | --- | --- | --- | --- |
| #1 | 60 to 690 ms | 190 to 990 ms | 0.0001 | \*\*\* | 1.8977 | positive |
| #2 | 970 to 1130 ms | 210 to 990 ms | 0.037 | \* | -1.5081 | negative |

  

all electrodes

|  | train times | test times | cluster p | cluster p-star | peak Cohen's d | direction |
| --- | --- | --- | --- | --- | --- | --- |
| #1 | 60 to 290 ms | 200 to 990 ms | 0.0026 | \*\* | 1.8401 | positive |
| #2 | 360 to 700 ms | 210 to 990 ms | 0.0058 | \*\* | 1.6888 | positive |
| #3 | 1220 to 1290 ms | 230 to 990 ms | 0.0371 | \* | 1.9323 | positive |

#### E) trained on familiar vs. remembered objects, tested on personally familiar faces

  

  

left anterior

|  | train times | test times | cluster p | cluster p-star | peak Cohen's d | direction |
| --- | --- | --- | --- | --- | --- | --- |
| #1 | 150 to 360 ms | 200 to 820 ms | 0.0473 | \* | 1.3243 | positive |
| #2 | 390 to 1260 ms | 220 to 990 ms | 0.0003 | \*\*\* | -1.0667 | negative |

  

right anterior

|  | train times | test times | cluster p | cluster p-star | peak Cohen's d | direction |
| --- | --- | --- | --- | --- | --- | --- |
| #1 | 150 to 420 ms | 200 to 990 ms | 0.0111 | \* | 1.4757 | positive |
| #2 | 440 to 1150 ms | 210 to 990 ms | 0.0003 | \*\*\* | -1.334 | negative |

  

left central

|  | train times | test times | cluster p | cluster p-star | peak Cohen's d | direction |
| --- | --- | --- | --- | --- | --- | --- |
| #1 | 20 to 560 ms | 170 to 990 ms | 0.0001 | \*\*\* | 1.8795 | positive |
| #3 | 1060 to 1290 ms | 180 to 990 ms | 0.0023 | \*\* | 1.7307 | positive |
| #4 | 530 to 820 ms | 370 to 870 ms | 0.0146 | \* | -1.7715 | negative |
| #2 | 850 to 1040 ms | 200 to 870 ms | 0.0402 | \* | 1.358 | positive |

  

right central

|  | train times | test times | cluster p | cluster p-star | peak Cohen's d | direction |
| --- | --- | --- | --- | --- | --- | --- |
| #1 | 20 to 530 ms | 180 to 990 ms | 0.0001 | \*\*\* | 1.9626 | positive |
| #3 | 510 to 1060 ms | 310 to 990 ms | 0.0001 | \*\*\* | -1.591 | negative |
| #2 | 1060 to 1290 ms | 200 to 990 ms | 0.0024 | \*\* | 1.8651 | positive |

  

left posterior

|  | train times | test times | cluster p | cluster p-star | peak Cohen's d | direction |
| --- | --- | --- | --- | --- | --- | --- |
| #1 | -20 to 470 ms | 170 to 820 ms | 0.0035 | \*\* | 1.2627 | positive |
| #3 | 480 to 910 ms | 380 to 990 ms | 0.0129 | \* | -0.7435 | negative |
| #2 | 1070 to 1290 ms | 180 to 810 ms | 0.0187 | \* | 1.5544 | positive |

  

right posterior

|  | train times | test times | cluster p | cluster p-star | peak Cohen's d | direction |
| --- | --- | --- | --- | --- | --- | --- |
| #1 | 170 to 470 ms | 200 to 990 ms | 0.0017 | \*\* | 1.4143 | positive |
| #3 | 490 to 1120 ms | 190 to 990 ms | 0.0014 | \*\* | -1.1567 | negative |
| #2 | 1220 to 1290 ms | 200 to 990 ms | 0.0363 | \* | 1.8866 | positive |

  

all electrodes

|  | train times | test times | cluster p | cluster p-star | peak Cohen's d | direction |
| --- | --- | --- | --- | --- | --- | --- |
| #1 | -20 to 590 ms | 150 to 990 ms | 0.0013 | \*\* | 1.8963 | positive |
| #2 | 500 to 910 ms | 460 to 880 ms | 0.0188 | \* | -0.7634 | negative |
