## Supplementary Table for "Shared neural codes of recognition memory": Supplementary_Table_5.html

  


### Supplementary Table 5.

  

Results of the main spatio-temporal searchlight analyses  

#### A) personally familiar and unfamiliar faces, leave-one-subject-out

  

|  | #1 |
| --- | --- |
| start time | 150.0 |
| stop time | 990.0 |
| peak time | 470.0 |
| peak channel | FT10 |
| cluster p | 0.0001 |
| peak Cohen's d | 3.021947 |
| direction | positive |

#### B) trained on experimentally familiarized face and unfamiliar faces, tested on personally familiar and unfamiliar faces

  

|  | #1 | #2 |
| --- | --- | --- |
| start time | 190 | 690 |
| stop time | 850 | 990 |
| peak time | 470 | 960 |
| peak channel | CP1 | P10 |
| cluster p | 0.0001 | 0.0234 |
| peak Cohen's d | 2.325809 | -1.731099 |
| direction | positive | negative |

#### C) trained on famous and unfamiliar faces, tested on personally familiar and unfamiliar faces

  

|  | #1 |
| --- | --- |
| start time | 190 |
| stop time | 990 |
| peak time | 550 |
| peak channel | PO8 |
| cluster p | 0.0001 |
| peak Cohen's d | 1.575951 |
| direction | positive |

#### D) trained on familiar and unfamiliar music, tested on personally familiar and unfamiliar faces

  

|  | #1 |
| --- | --- |
| start time | 270 |
| stop time | 990 |
| peak time | 470 |
| peak channel | CPz |
| cluster p | 0.0001 |
| peak Cohen's d | 1.727047 |
| direction | positive |

#### E) trained on remembered and forgotten object-scene associations, tested on personally familiar and unfamiliar faces

  

|  | #1 |
| --- | --- |
| start time | 200 |
| stop time | 990 |
| peak time | 480 |
| peak channel | Pz |
| cluster p | 0.0001 |
| peak Cohen's d | 1.858544 |
| direction | positive |

#### F) trained on familiar and novel objects, tested on personally familiar and unfamiliar faces

  

|  | #1 | #2 |
| --- | --- | --- |
| start time | 190 | 560 |
| stop time | 660 | 990 |
| peak time | 480 | 720 |
| peak channel | PO4 | P2 |
| cluster p | 0.0001 | 0.0046 |
| peak Cohen's d | 1.907891 | -1.351184 |
| direction | positive | negative |

#### G) trained on remembered and novel objects, tested on personally familiar and unfamiliar faces

  

|  | #1 |
| --- | --- |
| start time | 220 |
| stop time | 990 |
| peak time | 560 |
| peak channel | P8 |
| cluster p | 0.0001 |
| peak Cohen's d | 2.088025 |
| direction | positive |
