## Supplementary Table for "Shared neural codes of recognition memory": Supplementary_Table_6.html

  


### Supplementary Table 6.

  

Results of the spatio-temporal searchlight analyses, trained on the familiar - remembered object dataset  

#### A) trained on FA familiar and novel objects, tested on personally familiar and unfamiliar faces

  

|  | #1 |
| --- | --- |
| start time | 180 |
| stop time | 940 |
| peak time | 500 |
| peak channel | P4 |
| cluster p | 0.0001 |
| peak Cohen's d | 1.705797 |
| direction | positive |

#### B) trained on FA remembered and novel objects, tested on personally familiar and unfamiliar faces

  

|  | #1 |
| --- | --- |
| start time | 190 |
| stop time | 990 |
| peak time | 550 |
| peak channel | P8 |
| cluster p | 0.0001 |
| peak Cohen's d | 1.752525 |
| direction | positive |

#### C) trained on forgotten vs. novel objects, tested on personally familiar and unfamiliar faces

  

|  | #1 |
| --- | --- |
| start time | 160 |
| stop time | 990 |
| peak time | 500 |
| peak channel | P4 |
| cluster p | 0.0001 |
| peak Cohen's d | 1.852643 |
| direction | positive |

#### D) trained on objects, incorrect answers with subjective labels, tested on personally familiar and unfamiliar faces

  

|  | #1 | #2 |
| --- | --- | --- |
| start time | 190 | 840 |
| stop time | 920 | 990 |
| peak time | 580 | 930 |
| peak channel | PO7 | C1 |
| cluster p | 0.0001 | 0.0416 |
| peak Cohen's d | 1.820817 | -1.218869 |
| direction | positive | negative |

#### E) trained on familiar vs. remembered objects, tested on personally familiar faces

  

|  | #1 | #2 |
| --- | --- | --- |
| start time | 190 | 370 |
| stop time | 530 | 990 |
| peak time | 460 | 600 |
| peak channel | O2 | P8 |
| cluster p | 0.0022 | 0.0001 |
| peak Cohen's d | 1.742299 | -2.267768 |
| direction | familiar | remembered |
